# Chromatin state shapes site-specific A-to-I RNA editing

**DOI:** 10.64898/2026.07.27.740859

**Authors:** Peng Cai, Xing Chen, Jun Guo, Yuanmei Wang, Huanming Yang, Li Lin, Yi Zhang

## Abstract

Adenosine-to-inosine (A-to-I) RNA editing is a widespread post-transcriptional mechanism that diversifies the transcriptome. While ADAR enzymes catalyze this reaction, the upstream mechanisms that determine why individual adenosines are edited at markedly different efficiencies remain poorly understood. Here, we integrated multi-omics datasets from human cell lines and mouse embryonic tissues and developed machine learning models that distinguish high-and low-frequency editing sites based on local epigenetic features. Across species, tissues, and developmental stages, H3K36me3 consistently emerges as the strongest negative predictor of editing frequency, whereas the histone variant H2A.Z.1 shows a positive association. Functional validations in H2AFZ-knockdown cells reveal that H2A.Z.1 preferentially facilitates editing at high-frequency sites (76.3% of sites decreased, mean Δ =-0.07), whereas SETD2-knockout-mediated loss of H3K36me3 selectively derepresses editing at low-frequency sites (86.9% upregulated, ∼2.7-fold increase). These findings establish chromatin state as an upstream regulatory layer that modulates RNA editing independently of editing enzyme abundance and reveal opposing roles for H3K36me3 and H2A.Z.1 in shaping RNA editing landscapes. Together, our study provides a conceptual framework linking epigenetic regulation to post-transcriptional RNA modification, suggesting that chromatin-mediated regulation contributes to the establishment of site-specific RNA editing programs across mammalian genomes.

## INTRODUCTION

RNA editing is a post-transcriptional regulatory mechanism that expands transcriptomic and proteomic diversity. Genome-wide studies have identified widespread differentially edited loci across diverse physiological and pathological conditions, highlighting the broad functional significance of RNA editing [1]. Among the various forms of RNA editing, A-to-I editing, catalyzed by the ADAR family of enzymes [2–6], is the predominant type in animals. By recoding adenosines into inosines, which are interpreted as guanosines during translation and RNA processing [7, 8], this biochemical conversion can alter codon identity, modulate alternative splicing, reshape RNA secondary structure, and rewire RNA-protein and RNA-RNA interactions [9–11]. As such, ADAR-mediated editing is essential for suppressing aberrant innate immune activation and maintaining immune homeostasis [12, 13]. Dysregulated RNA editing has been implicated in multiple human diseases, including cancer [14], autoimmune disorders [15], and neurological diseases such as amyotrophic lateral sclerosis [16], epilepsy [17], depression [18], schizophrenia [19], and bipolar disorder [19]. Therefore, these findings underscore the biological and clinical importance of understanding the mechanisms regulating RNA editing [20].

Although ADARs catalyze RNA editing directly, ADAR abundance alone cannot explain the remarkable variability in editing efficiency observed among individual editing sites, cell types, developmental stages, and physiological conditions [21]. Previous studies have identified several regulatory factors, including local RNA sequence context, RNA secondary structure, RNA-binding proteins, ADAR-interacting proteins, and RNA modifications such as m6A [22–31]. Yet these factors do not fully account for the extensive heterogeneity observed across the transcriptome.

A fundamental question remains: does the chromatin state of the DNA template from which RNA is transcribed predetermine which sites become edited? This question is motivated by two observations. First, co-transcriptional RNA processing, including alternative splicing, polyadenylation, and m6A modification, is increasingly recognized as shaped by chromatin architecture [32–37]. Second, A-to-I RNA editing occurs predominantly co-transcriptionally [38], and ADAR substrates (double-stranded RNA structures) form during transcription [39], suggesting that transcriptional dynamics can affect editing efficiency. However, whether histone modifications actively modulate RNA editing has not been systematically investigated.

Here, we hypothesize that chromatin state represents an upstream regulatory layer controlling A-to-I RNA editing. To test this hypothesis, we developed an integrative framework and systematically evaluated the contribution of epigenetic features to RNA editing frequency across species and developmental contexts. We further leveraged knockout and knockdown studies to functionally validate the role of prominent epigenetic marks in regulating RNA editing.

## RESULTS

### Characterization of A-to-I RNA editing events in human cell lines

To systematically identify A-to-I RNA editing events, we established a computational pipeline (Figure 1A). Briefly, raw RNA-seq reads were subjected to stringent quality control to remove low-quality sequences and adapter contamination, followed by alignment to the reference genome. A-to-I RNA editing sites were subsequently identified using REDItools [40].

**Figure 1.**
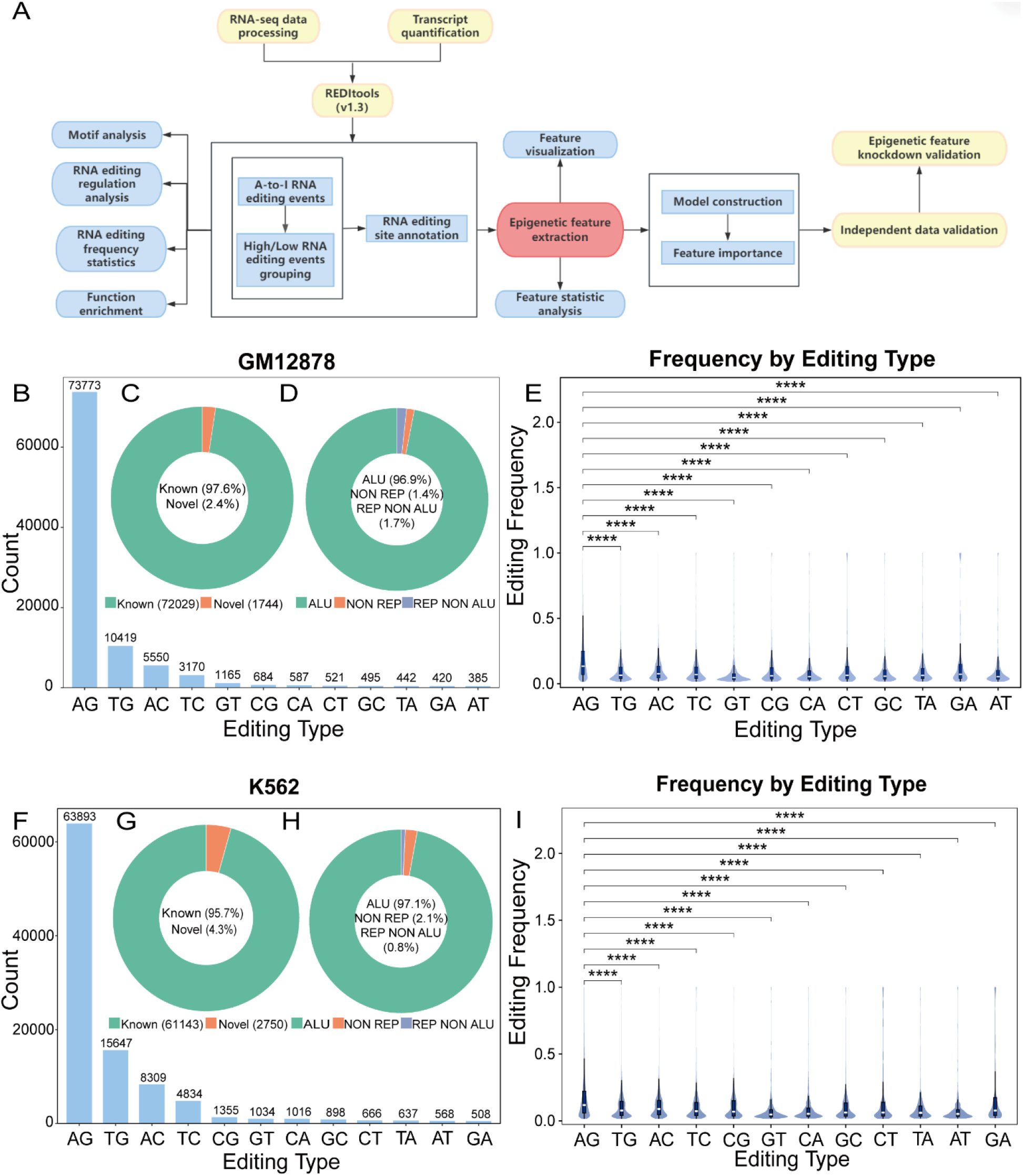
Identification and characterization of A-to-I RNA editing in human cell lines. (A) Schematic of the computational pipeline for RNA editing analysis. (B–D) Summary statistics of editing sites identified in GM12878 cells. (B) Total number of sites in each editing type. (C) Editing sites categorized by annotation status. (D) Proportion of sites located within Alu repetitive elements versus non-repetitive regions. (E) Relative frequencies of all 12 nucleotide substitution types in GM12878, illustrating the predominant A-to-G signature characteristic of A-to-I editing, ****p < 0.0001. (F–H) Summary statistics for editing sites identified in K562 cells. (F) Total number of sites in each editing type. (G) Annotation status of editing sites. (H) Proportion of sites located within Alu repetitive elements versus non-repetitive regions. (I) Relative frequencies of nucleotide substitution types in K562, confirming A-to-G as the predominant editing signature, ****p < 0.0001.

We first applied this pipeline to RNA-seq datasets from the GM12878, K562, and HeLa-S3 human cell lines generated by the ENCODE project [41, 42]. In GM12878 cells, we identified 73,773 A-to-I RNA editing sites, of which 97.6% overlapped with previously annotated sites, while 1,744 were putative novel. Consistent with the established preference of ADAR enzymes for double-stranded RNA formed by inverted Alu repeats [43], 96.9% of these editing sites were located within Alu repetitive elements (Figures 1B–D). Moreover, A-to-G substitutions overwhelmingly predominated over all other nucleotide substitutions, confirming the characteristic signature of A-to-I RNA editing (Figure 1E). Analysis of K562 cells identified 63,893 A-to-I editing sites, including 61,143 annotated and 2,750 putative novel sites. Similar to GM12878 cells, editing events were highly enriched within Alu elements (97.1%) and exhibited nearly identical genomic distribution patterns (Figures 1F–I). HeLa-S3 cells displayed comparable editing profiles (Supplementary Figure 1A-B), demonstrating that the overall landscape of A-to-I RNA editing is highly conserved across diverse human cell types.

To define the local sequence features associated with A-to-I editing, we examined nucleotide composition within ±2 nucleotides surrounding each editing site. Reverse-complement sequences were used for editing sites located on the negative strand to maintain a uniform transcriptional orientation. Motif analysis using MEME-ChIP identified a conserved sequence signature characterized by depletion of guanosine immediately upstream (−1 position) and enrichment immediately downstream (+1 position) of the edited adenosine (Figure 2A-B and Supplementary Figure 1D). This motif was remarkably consistent across GM12878, K562, and HeLa-S3 cells (Figure 2C-D and Supplementary Figure 1E), in agreement with previously reported ADAR editing preferences [22, 23].

**Figure 2.**
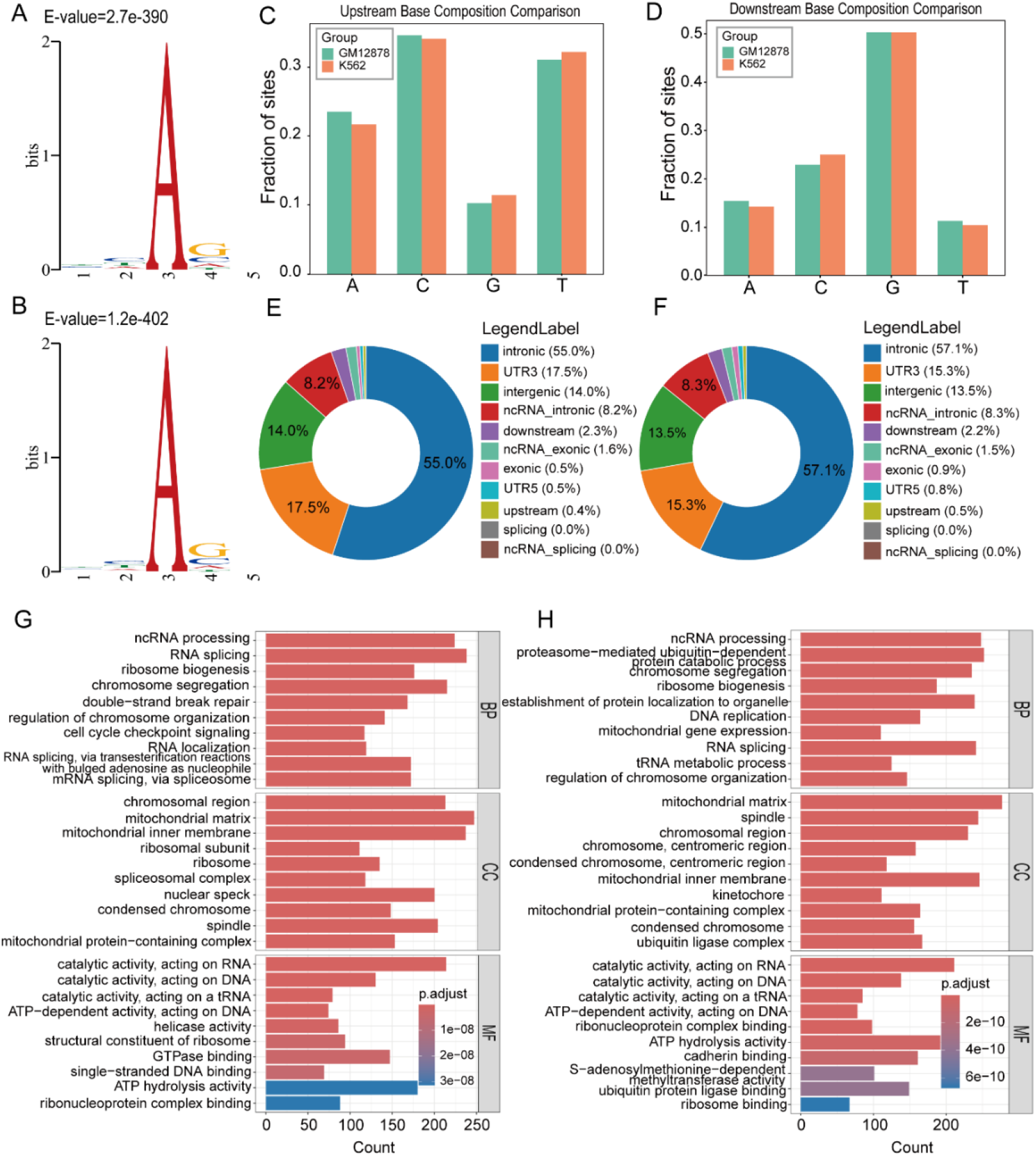
Genomic distribution and functional enrichment of A-to-I RNA editing sites in human cell lines. (A-B) Sequence logos showing the 2-bp nucleotide composition immediately upstream and downstream of editing sites in (A) GM12878 and (B) K562 cells. (C) The 1 bp nucleotide composition at upstream (-1) of RNA editing sites in both cell lines. (D) The 1 bp nucleotide composition at downstream (+1) of RNA editing sites in both cell lines. (E) Genomic distribution of RNA editing sites in the GM12878 cell line. (F) Genomic distribution of RNA editing sites in the K562 cell line. (G) Gene Ontology (GO) enrichment analysis of host genes harboring editing sites in GM12878 cells. (H) Gene Ontology (GO) enrichment analysis of host genes harboring editing sites in K562 cells. BP, biological process; MF, molecular function; CC, cellular component.

We next examined the genomic distribution of editing sites using ANNOVAR-based annotation [44]. In both GM12878 and K562 cells, the majority of editing sites were located within intronic regions (> 50%), followed by substantial enrichment in 3′ untranslated regions (3’UTRs) (Figures 2E-F). A similar distribution was observed in HeLa-S3 cells, where 45.6% and 24.6% of editing sites were mapped to introns and 3’UTRs, respectively (Supplementary Figure 1F).

To investigate the potential biological functions of edited transcripts, we mapped editing sites to their host genes and performed Gene Ontology (GO) enrichment analysis. Edited genes were significantly enriched in biological processes related to RNA metabolism, including RNA processing, RNA splicing, ribosome biogenesis, and chromosome segregation. Cellular component analysis highlighted enrichment in the mitochondrial matrix and chromosomal regions, whereas molecular function analysis revealed overrepresentation of catalytic activities acting on RNA and DNA, as well as ATP hydrolysis activity (Figures 2G-H and Supplementary Figure 1C).

Collectively, these results demonstrate that A-to-I RNA editing is widespread and highly conserved across human cell lines. The predominance of editing within Alu repetitive elements, together with the conserved sequence motifs and genomic distributions, is consistent with previous reports and validates the robustness and accuracy of our RNA editing analysis pipeline [43, 45, 46].

### Correlation of ADAR expression and chromatin features with A-to-I RNA editing frequency in human cell lines

ADAR1 is ubiquitously expressed across human tissues [2], whereas ADAR2 is most abundant in the brain but is also expressed at lower levels in other tissues [4]. Although both enzymes catalyze A-to-I RNA editing, they exhibit distinct substrate preferences: ADAR1 predominantly edits repetitive elements, whereas ADAR2 preferentially targets non-repetitive coding regions [21].

To identify factors associated with RNA editing activity, we first quantified the expression of ADAR family members in GM12878, K562, and HeLa-S3 cells using RNA-seq data. Consistent with previous studies [47], ADAR1 was the predominant editing enzyme in all three cell lines, whereas ADAR2 was expressed at substantially lower levels (Figure 3A). In contrast, ADAR3, which lacks deaminase activity, was barely detectable. We next calculated the global RNA editing frequency for each sample, defined as the proportion of sequencing reads supporting A-to-G substitutions across all identified editing sites. ADAR1 expression exhibited a strong positive correlation with global RNA editing frequency (R² = 0.87), whereas ADAR2 expression showed no significant association (Figure 3B-C). These findings indicate that ADAR1 is the principal determinant of transcriptome-wide A-to-I RNA editing in these cell lines, whereas it doesn’t explain site-specific variation of RNA editing.

**Figure 3.**
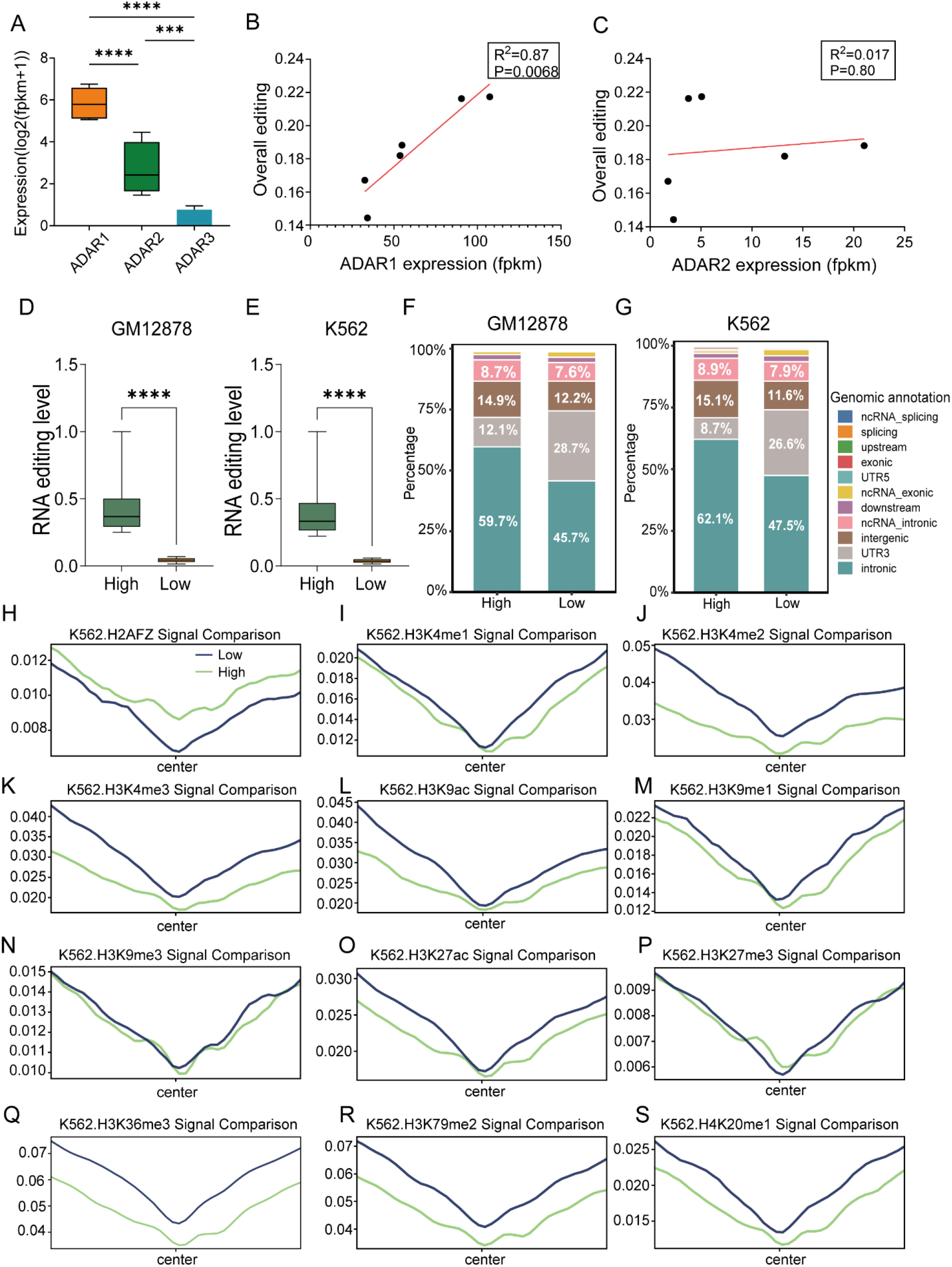
Correlation analysis for ADAR expression and epigenetic marks related to RNA editing frequency. (A) ADAR1, ADAR2, and ADAR3 transcript levels (FPKM) in GM12878, K562, and HeLa-S3 cells. (B) Correlation between ADAR1 expression and global A-to-I RNA editing frequency (R² = 0.87). (C) Correlation between ADAR2 expression and global editing frequency. (D, E) Number of editing sites classified as high-frequency (≥75th percentile) or low-frequency (≤25th percentile) in (D) GM12878 and (E) K562. (F, G) Genomic distribution of high-frequency and low-frequency editing sites in (F) GM12878 and (G) K562. (H–S) Metagene plots showing average ChIP-seq signal intensity within ±150 bp of high-frequency (green; ≥75th percentile) and low-frequency (blue; ≤25th percentile) editing sites in K562 cells for (H) H2A.Z.1, (I) H3K4me1, (J) H3K4me2, (K) H3K4me3, (L) H3K9ac, (M) H3K9me1, (N) H3K9me3, (O) H3K27ac, (P) H3K27me3, (Q) H3K36me3, (R) H3K79me2, and (S) H4K20me1. ***p < 0.001,****p < 0.0001.

Despite this strong global correlation, the expression levels of ADAR1 and ADAR2 could not account for the marked heterogeneity in editing frequencies among individual sites, suggesting that local regulatory mechanisms contribute to site-specific editing efficiency [21]. To investigate this possibility, editing sites were ranked according to editing frequency and classified into high-frequency (≥75th percentile) and low-frequency (≤25th percentile) groups (Figure 3D-E). Comparison of their genomic distributions revealed a significant shift in genomic localization: the distribution of editing sites within 3’UTRs is significantly higher in the low-frequency group compared to the high-frequency group; whereas more intronic editing sites were mapped for the high-frequency group than the low-frequency group (Figure 3F-G). The enrichment of low-frequency sites in 3’UTRs and high-frequency sites in introns suggests that genomic context influences editing efficiency, potentially through differential chromatin environments associated with these regions.

We next examined whether local chromatin states contribute to this site-specific variability. If chromatin architecture influences RNA editing, high-and low-frequency editing sites would be expected to exhibit distinct epigenetic signatures. To test this hypothesis, we compared histone modification signals within ±150 bp of editing sites between the two groups. In K562 cells, high-frequency editing sites displayed a distinct chromatin landscape. H2A.Z.1 (encoded by H2AFZ), a histone variant associated with nucleosome instability and transcriptional plasticity [48], was significantly enriched around high-frequency editing sites, suggesting a more permissive chromatin environment (Figure 3H). In contrast, the heterochromatin marks H3K27me3 and H3K9me3 showed little difference between the two groups. Notably, several histone modifications associated with actively transcribed gene bodies or promoters, including H3K36me3, H3K79me2, H3K4me3, and H3K4me2, exhibited significantly lower signal intensity surrounding high-frequency editing sites than low-frequency sites (Figures 3I–S and Table 1). Similar chromatin patterns were independently observed in GM12878 cells (Supplementary Figure 2 and Supplementary Table 1), indicating that these epigenetic associations are reproducible across cell types. Among all histone features examined, H2A.Z.1 consistently emerged as the strongest positive correlate of high-frequency RNA editing.

**Table 1.** Statistical comparison of ChIP-seq signals for each histone mark within ±150 bp of editing sites between high-and low-frequency editing groups in K562 cells, p-values were adjusted using the Benjamini–Hochberg (BH) procedure.

| Histone mark | High_frequency<br>mean | Low_frequency<br>mean | Fold change<br>(High/Low) | BH-FDR<br>q-value |
| --- | --- | --- | --- | --- |
| H3K36me3 | 13.22 | 16.97 | 0.78 | 7.3e-155 |
| H4K20me1 | 4.62 | 5.58 | 0.83 | 5.5e-42 |
| H3K79me2 | 12.62 | 15.58 | 0.81 | 5.7e-18 |
| H3K27ac | 6.03 | 6.72 | 0.90 | 5.4e-07 |
| H3K9me1 | 4.77 | 5.23 | 0.91 | 1.8e-19 |
| H3K9ac | 6.83 | 8.14 | 0.84 | 1.3e-06 |
| H3K4me1 | 4.17 | 4.56 | 0.92 | 4.8e-11 |
| H3K4me3 | 6.43 | 8.31 | 0.77 | 3.8e-13 |
| H3K4me2 | 7.65 | 10.06 | 0.76 | 1.9e-12 |
| H3K9me3 | 3.51 | 3.61 | 0.97 | 0.0139 |
| H3K27me3 | 2.16 | 2.19 | 0.98 | 0.054 |
| H2A.Z.1 | 2.95 | 2.58 | 1.14 | 2.8e-05 |

Therefore, these findings demonstrated that while ADAR1 expression is the primary determinant of global A-to-I RNA editing activity, local chromatin architecture may contribute substantially to site-specific editing variability. In particular, enrichment of H2A.Z.1 and depletion of several active histone marks distinguish highly edited sites, suggesting that the epigenetic landscape is likely an important regulator of RNA editing efficiency.

### Model prediction and feature importance analysis for A-to-I RNA editing frequency in human cell lines

To determine whether local epigenetic features can predict RNA editing efficiency, we developed binary classification models to distinguish high-and low-frequency A-to-I RNA editing sites using histone mark signals within ±150 bp of each editing site in GM12878 and K562 cells. We benchmarked nine classification frameworks: Logistic Regression, Support Vector Machine, Random Forest, eXtreme Gradient Boosting (XGBoost), Naïve Bayes, CatBoost, ResNet-style MLP, Extremely Randomized Trees (Extra Trees) classifier, and Extra Trees Full Ratio Regressor. All models were evaluated with identical stratified five-fold partitions; performance was quantified by the area under the receiver operating characteristic curve (AUROC) and precision-recall curve (AUPRC). Among these models, Random Forest and Extra Trees consistently achieved the highest predictive performance, with five-fold cross-validation AUROC and AUPRC values of approximately 0.70 and 0.69, respectively, across both cell lines (Table 2 and Figure 4A–B). Given the dominant role of ADAR enzymes in A-to-I RNA editing, this level of predictive accuracy is impressive, indicating that local chromatin features carry important informative signals associated with site-specific editing frequency.

**Figure 4.**
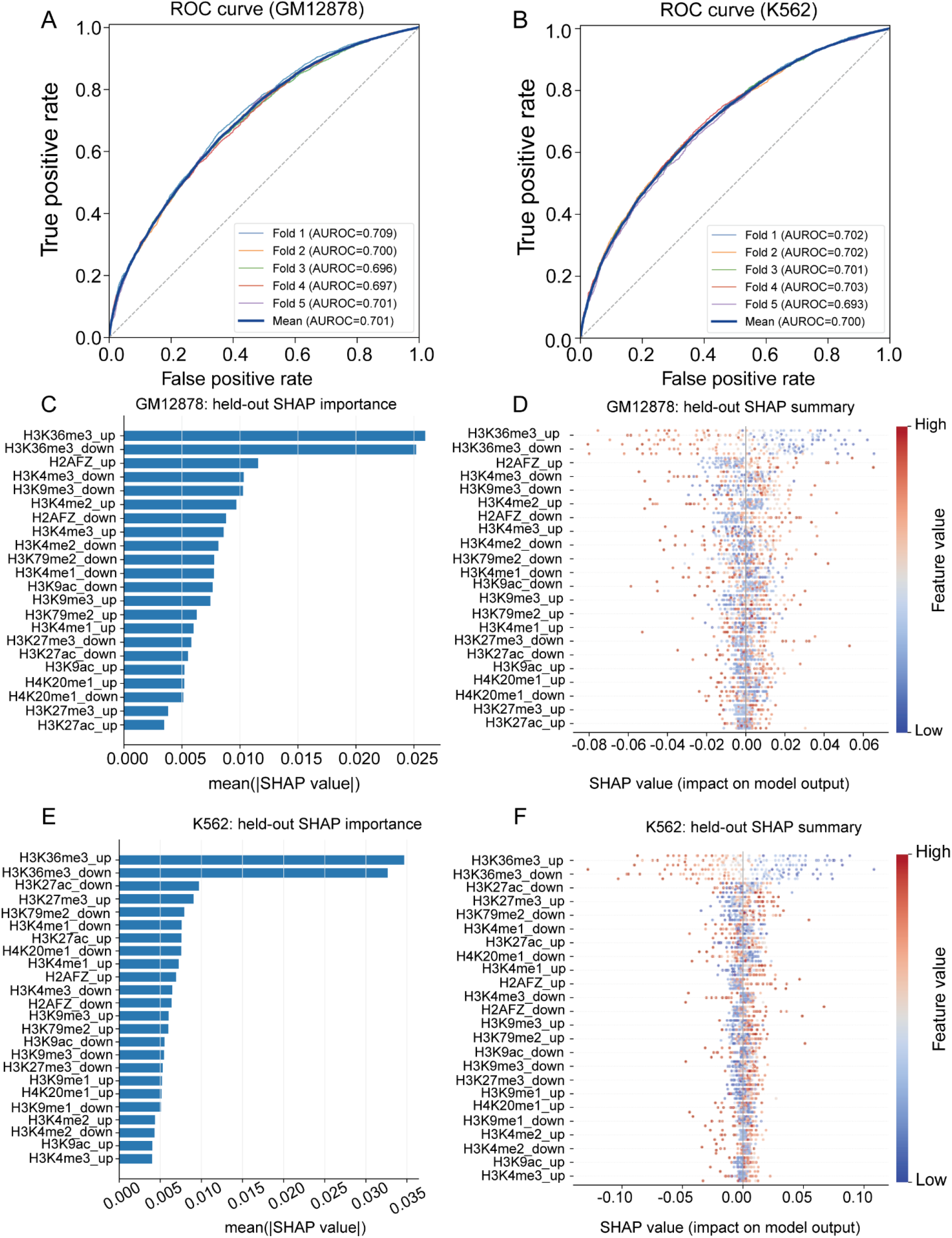
Machine learning prediction of editing frequency and model interpretation. (A) AUROC curve of the Random Forest binary classifier for high-versus low-frequency editing sites in GM12878 (five-fold cross-validation). (B) AUROC curve for the corresponding model in K562. (C) SHAP feature importance bar plot for the GM12878 model. (D) SHAP summary plot showing the distribution of feature contributions in GM12878. (E) SHAP feature importance bar plot for the K562 model. (F) SHAP summary plot showing the distribution of feature contributions in K562.

**Table 2.** Model performance for predicting high-and low-frequency A-to-I RNA editing groups using histone mark signals within ±150 bp of each editing site in GM12878 and K562 cells.

| Model | GM12878 |  | K562 |  |
| --- | --- | --- | --- | --- |
|  | AUROC | AUPRC | AUROC | AUPRC |
| Logistic Regression | 0.5713 | 0.5672 | 0.6039 | 0.5882 |
| Support Vector Machine | 0.5930 | 0.5844 | 0.6212 | 0.6128 |
| Random Forest | 0.7005 | 0.6978 | 0.7003 | 0.6933 |
| XGBoost | 0.5965 | 0.5937 | 0.6238 | 0.6150 |
| Naïve Bayes | 0.5576 | 0.5566 | 0.5731 | 0.5578 |
| CatBoost | 0.6086 | 0.6063 | 0.6242 | 0.6166 |
| ResNet-style MLP | 0.5903 | 0.5870 | 0.6159 | 0.6069 |
| Extra Trees classifier | 0.7070 | 0.7047 | 0.7066 | 0.6992 |
| Extra Trees Full Ratio Regressor | 0.7322 | 0.7188 | 0.7453 | 0.7269 |

Because RNA editing frequencies are inherently ratio-scale variables (ranging from 0 to 1, representing the proportion of edited transcripts), we additionally employed the Extra Trees Full Ratio Regressor [49]. This approach preserves the continuous structure by training on sites spanning the complete editing-ratio distribution while retaining their measured labels. This regression-based approach achieved an AUROC of 0.7453 and an AUPRC of 0.7269 in K562 cells, outperforming all binary classifiers. Similar performance was observed for GM12878 as well (Table 2). By preserving the progression from low through intermediate to high editing frequencies, the regression model learns the relative ordering of sites more effectively, directly benefiting the ranking-based AUROC and AUPRC metrics.

To identify the epigenetic features driving model predictions, we performed SHapley Additive exPlanations (SHAP) analysis, which quantifies the contribution of individual features while accounting for feature interactions [50]. H3K36me3 was consistently ranked as the most influential predictor in both GM12878 and K562 cells (Figure 4C and E). Importantly, higher feature values of H3K36me3 in both up-and downstream of editing sites were associated with more negative SHAP values, indicating that elevated H3K36me3 levels negatively contribute to the model output (Figure 4D and F). This result suggests that, although multiple histone modifications contribute to editing-site classification, variation in H3K36me3 provides the greatest predictive information for distinguishing highly edited from weakly edited sites.

Therefore, these findings demonstrated that local chromatin features encode substantial information regarding RNA editing efficiency and can be leveraged to predict editing frequency at individual sites, supporting a model in which chromatin architecture contributes to the regulation of RNA editing beyond the effects of ADAR expression alone.

### Functional validation of H2A.Z.1 in regulating A-to-I RNA editing

H2A.Z is an essential, evolutionarily conserved histone H2A variant that is dynamically loaded and removed from nucleosomes, enabling rapid transcriptional responses to developmental and environmental cues [51]. Through modulation of nucleosome stability, H2A.Z regulates diverse chromatin-associated processes, including transcription, DNA repair, DNA replication, and development [48]. To date, three H2A.Z isoforms have been identified. H2A.Z.1 and H2A.Z.2 are encoded by two nonallelic genes H2AFZ and H2AFV, respectively, with the H2A.Z.1 being more abundantly expressed [52]. A third isoform, H2A.Z.2.2, is a splice variant of H2A.Z.2 [53].

Our correlative analyses identified H2A.Z.1 as the histone feature most positively associated with high-frequency A-to-I RNA editing. To determine whether H2A.Z.1 functionally contributes to RNA editing, we analyzed the H2AFZ knockdown RNA-seq dataset generated from the human epithelial cell line MCF-10A [54]. We first examined whether H2AFZ depletion altered the expression of the major RNA editing enzymes. RNA-seq analysis revealed that knockdown of H2AFZ did not significantly affect the expression of either ADAR1 or ADAR2 (Figure 5A-B), indicating that any changes in RNA editing are unlikely to result from altered ADAR abundance and instead reflect chromatin-dependent regulatory mechanisms.

**Figure 5.**
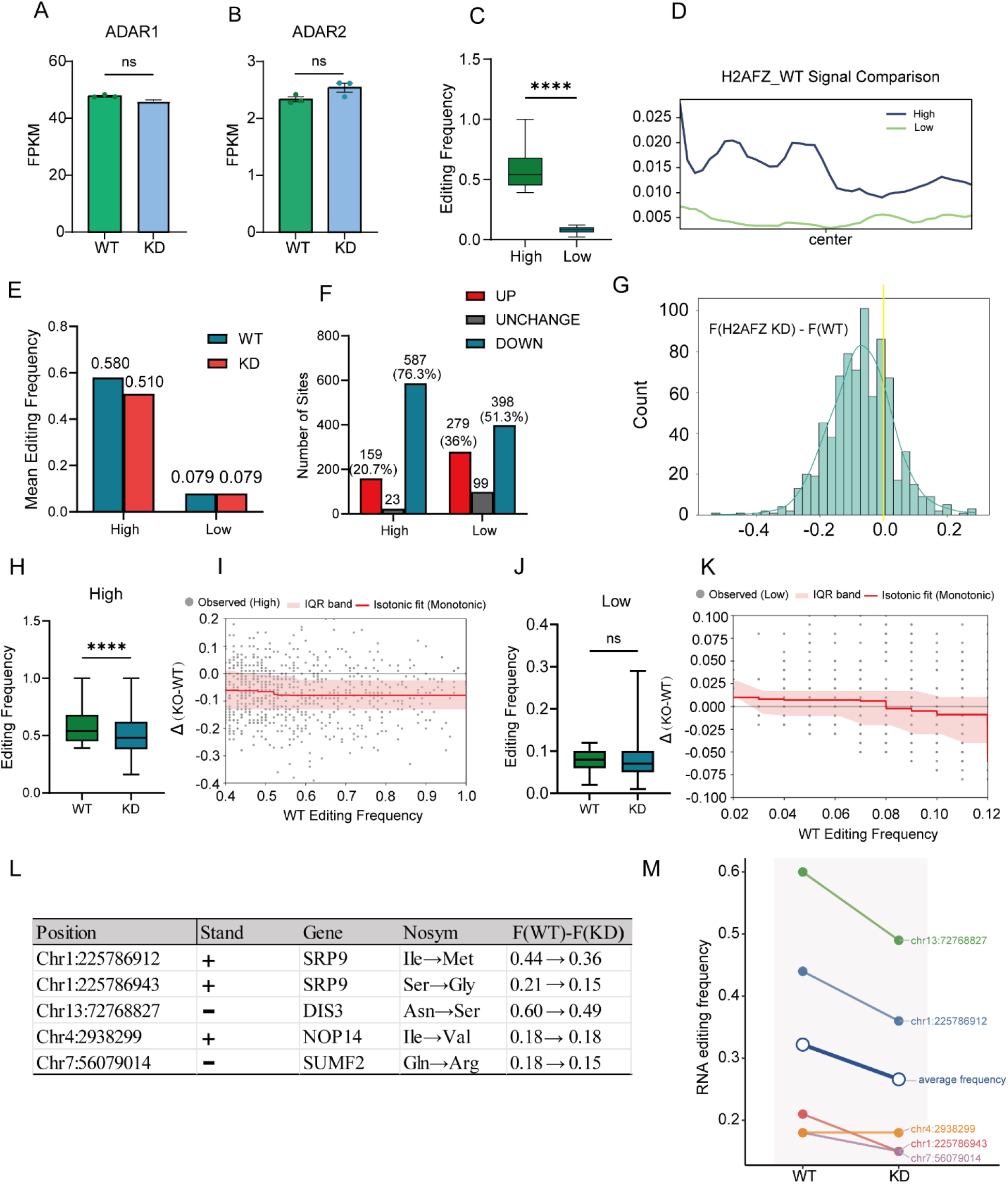
H2A.Z depletion selectively reduces editing at high-frequency A-to-I RNA editing sites. (A) ADAR1 transcript expression in wild-type (WT) and H2AFZ knockdown (KD) MCF-10A cells. (B) ADAR2 transcript expression in WT and H2AFZ KD cells. (C) Editing frequency distribution in WT cells, stratified into high-frequency (≥75th percentile) and low-frequency (≤25th percentile) groups. (D) H2A.Z.1 ChIP-seq signal intensity within ±150 bp of high-and low-frequency editing sites in WT cells. (E) Mean editing frequency of high-and low-frequency groups in WT and H2AFZ KD. (F) Proportion of high-frequency editing sites or low-frequency editing sites exhibiting increased, decreased, or unchanged editing following H2AFZ KD. (G) Distribution of editing frequency changes (Δ) for high-frequency sites upon H2AFZ KD. (H) Editing frequency comparison for the high-frequency group between WT and H2AFZ KD. (I) Isotonic regression fit of the change in editing frequency (KD − WT, y-axis) as a function of wild-type editing frequency (x-axis) for high-frequency sites. (J) Editing frequency comparison for the low-frequency group between WT and H2AFZ KD. (K) Isotonic regression analysis for the low-frequency group. (L) Editing frequency changes at individual nonsynonymous editing sites following H2AFZ KD. (M) Summary of nonsynonymous editing site responses following H2AFZ KD. ****p < 0.0001; ns, not significant (p > 0.05).

We next identified A-to-I RNA editing sites in wild-type and H2AFZ knockdown samples and compared editing frequencies between matched conditions. Editing sites from wild-type were classified into high-and low-frequency groups (≥ 75th percentile and ≤ 25th percentile, respectively) (Figure 5C), which exhibited distinct H2A.Z.1 enrichment profiles within ±150 bp of editing sites (Figure 5D). Consistent with our genome-wide analyses, high-frequency editing sites in wild-type cells displayed significantly greater H2A.Z.1 occupancy than low-frequency sites.

Following H2AFZ knockdown, the overall editing frequency of the high-frequency group decreased from 0.580 to 0.510, corresponding to a mean reduction of 0.07 (Figure 5E). In contrast, the average editing frequency of the low-frequency group remained essentially unchanged. Consistent with this overall reduction, 587 of 769 (76.3%) high-frequency editing sites exhibited decreased editing frequencies, whereas only 159 sites (20.7%) showed increased editing (Figure 5F). By comparison, the low-frequency group displayed substantially weaker effects, with 398 of 776 (51.3%) sites exhibiting decreased editing and 279 sites (36.0%) exhibiting increased editing.

The distribution of editing frequency changes further demonstrated a pronounced leftward shift for high-frequency editing sites following H2AFZ depletion (Figure 5G), indicating an overall reduction in editing frequency. Statistical analysis confirmed that editing frequencies were significantly lower in H2AFZ knockdown samples than in wild-type controls for the high-frequency group (Figure 5H). Consistent with this observation, isotonic regression of the editing change (KD – WT) against baseline editing frequency revealed a monotone decreasing relationship, indicating that sites with higher wild-type editing frequencies experienced the largest reductions following H2AFZ depletion (Figure 5I). In contrast, no significant difference was observed between wild-type and H2AFZ knockdown samples for the low-frequency group (Figure 5J), and isotonic regression of the editing change against baseline frequency showed no consistent directional trend for the low-frequency group (Figure 5K).

Finally, we examined nonsynonymous editing events, which have the potential to alter protein-coding sequences. Following H2AFZ knockdown, four of the five detected nonsynonymous editing sites exhibited increased editing frequencies, whereas one remaining site was unchanged (Figure 5L-M), suggesting that H2A.Z.1-mediated regulation may extend to functionally relevant coding regions.

Together, these findings provide functional evidence that H2A.Z.1 positively regulates high-frequency A-to-I RNA editing sites. Depletion of H2AFZ selectively reduced editing at the majority of highly edited loci while exerting minimal effects on low-frequency sites. Importantly, because ADAR1 and ADAR2 expression remained unchanged following H2AFZ knockdown, these results support a model in which H2A.Z.1 regulates RNA editing through modulation of the local chromatin environment rather than through transcriptional control of the editing enzymes.

### Characteristics of A-to-I RNA editing during mouse embryonic development

To determine whether the molecular features of A-to-I RNA editing are evolutionarily conserved, we analyzed RNA-seq datasets from five mouse tissues (forebrain, midbrain, hindbrain, heart, and liver) spanning multiple embryonic developmental stages [41]. Raw sequencing data were obtained from the ENCODE database, with at least two biological replicates per condition. Following stringent quality control to remove low-quality reads and adapter sequences, cleaned reads were aligned to the mouse reference genome using STAR, and A-to-I RNA editing sites were identified using REDItools. High-confidence editing sites were subsequently defined through rigorous filtering (Table 3).

**Table 3.** Counts of A-to-I RNA editing events in multiple mouse tissues across embryonic developmental stages.

| Tissue | e11 | e12 | e13 | e14 | e15 | e16 | p0 |
| --- | --- | --- | --- | --- | --- | --- | --- |
| Forebrain | 527 | 1333 | 1231 | 761 | 1209 | 999 | 1007 |
| Midbrain | 1063 | 1183 | 1208 | 818 | 1288 | 1156 | 1045 |
| Hindbrain | 1057 | 1198 | 1190 | 872 | 1213 | 1143 | 1112 |
| Heart | 842 | 1072 | 1035 | 710 | 998 | 854 | 1016 |
| Liver | 912 | 1052 | 1058 | 641 | 814 | 871 | 952 |

Comparative analysis revealed that neural tissues (forebrain, midbrain, and hindbrain) generally harbored a greater number of RNA editing sites than non-neural tissues (heart and liver) across developmental stages (Figure 6A and Table 3). Despite these differences in site numbers, the global RNA editing frequency remained relatively stable at each developmental time point acrosstissues (Figure 6B), suggesting that overall editing activity is tightly regulated during embryogenesis.

**Figure 6.**
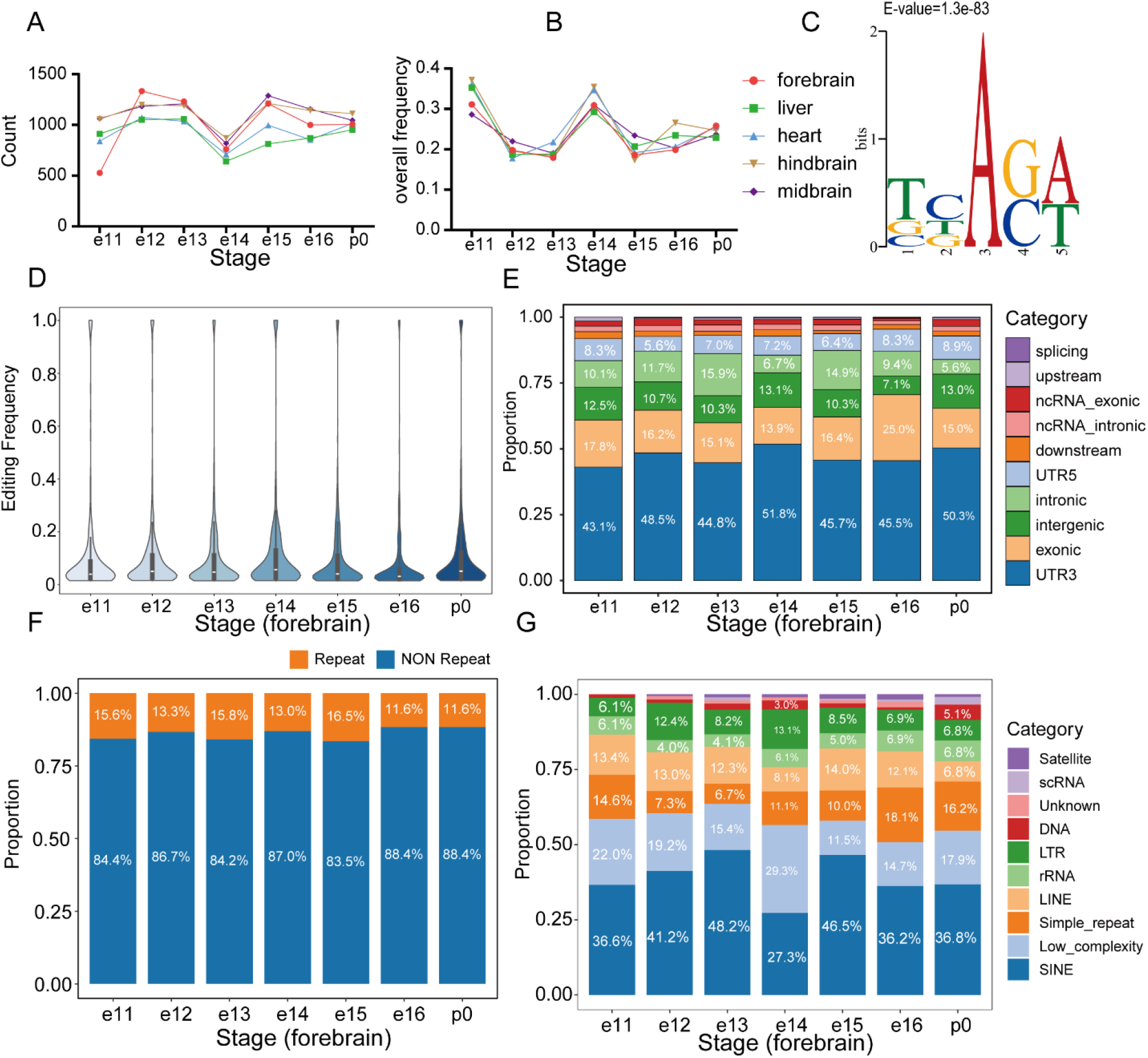
A-to-I RNA editing during mouse embryonic development. (A) Number of A-to-I editing sites identified in forebrain, midbrain, hindbrain, heart, and liver from embryonic day 11 (e11) to postnatal day 0 (p0). (B) Global A-to-I editing frequency across the five tissues and developmental stages. (C) Nucleotide composition of 2-bp sequence motif flanking A-to-I editing sites in the mouse forebrain. (D) Distribution of A-to-I editing frequencies in the mouse forebrain across developmental stages (e11–p0). (E) Genomic distribution of A-to-I editing sites in the mouse forebrain. (F) Proportion of A-to-I editing sites in repetitive versus non-repetitive elements in the mouse forebrain. (G) Genomic distribution of A-to-I editing sites within repetitive elements in the mouse forebrain.

We next characterized the sequence and genomic features of editing sites across developmental stages. Motif analysis identified a highly conserved sequence preference, with depletion of guanosine immediately upstream (−1 position) and enrichment immediately downstream (+1 position) of the edited adenosine (Figure 6C and Supplementary Figures 4-7). This signature is consistent with the canonical substrate preference of ADAR enzymes and closely recapitulates the motif observed in human cell lines, highlighting strong evolutionary conservation of sequence determinants governing A-to-I RNA editing. Furthermore, the distribution of editing frequencies remained highly consistent throughout embryonic development (Figure 6D and Supplementary Figures 4-7), underscoring the robustness of RNA editing regulation across developmental stages.

Analysis of genomic context revealed notable species-specific differences in editing site distribution. Unlike human cells, where editing events are predominantly enriched within intronic and Alu elements, mouse editing sites were primarily localized to 3’UTRs (Figure 6E and Supplementary Figures 3-6). Consistent with the absence of primate-specific Alu repeats in the mouse genome, most editing sites were located within non-repetitive regions (Figure 6F and Supplementary Figures 3-6). Among repetitive sequences, editing events were preferentially enriched in SINE elements (Figure 6G and Supplementary Figures 3-6), suggesting that species-specific repeat architectures contribute to distinct genomic landscapes of A-to-I RNA editing.

Thus, these results demonstrated that while the genomic distribution of RNA editing sites differs between species, key features, including sequence motifs and global editing activity, are highly conserved. The enrichment of editing sites in neural tissues further highlights a potential role for A-to-I RNA editing in brain development and function.

### Model prediction and feature importance analysis for A-to-I RNA editing frequency during mouse embryonic development

To investigate whether the epigenetic regulation of A-to-I RNA editing is conserved across species, we extended our analysis to mouse embryonic development, a period characterized by extensive chromatin remodeling and dynamic transcriptome regulation. A-to-I RNA editing sites identified across multiple tissues and developmental stages were stratified into high-and low-frequency groups according to their editing levels. We then developed a Random Forest–based binary classification model using local epigenetic features, including chromatin accessibility (ATAC-seq), eight histone modifications (H3K27ac, H3K27me3, H3K36me3, H3K4me1, H3K4me2, H3K4me3, H3K9ac, and H3K9me3), and DNA methylation levels within ±150 bp of each editing site. Model interpretation was performed using SHapley Additive exPlanations (SHAP) analysis to quantify the contribution of individual epigenetic features.

Across all five tissues and seven developmental stages, the model achieved stable and robust predictive performance, with AUC values consistently approaching 0.75 (Figure 7A). These results indicate that local epigenetic features contain substantial predictive information for RNA editing frequency during mouse embryogenesis, consistent with our observations in human cell lines.

**Figure 7.**
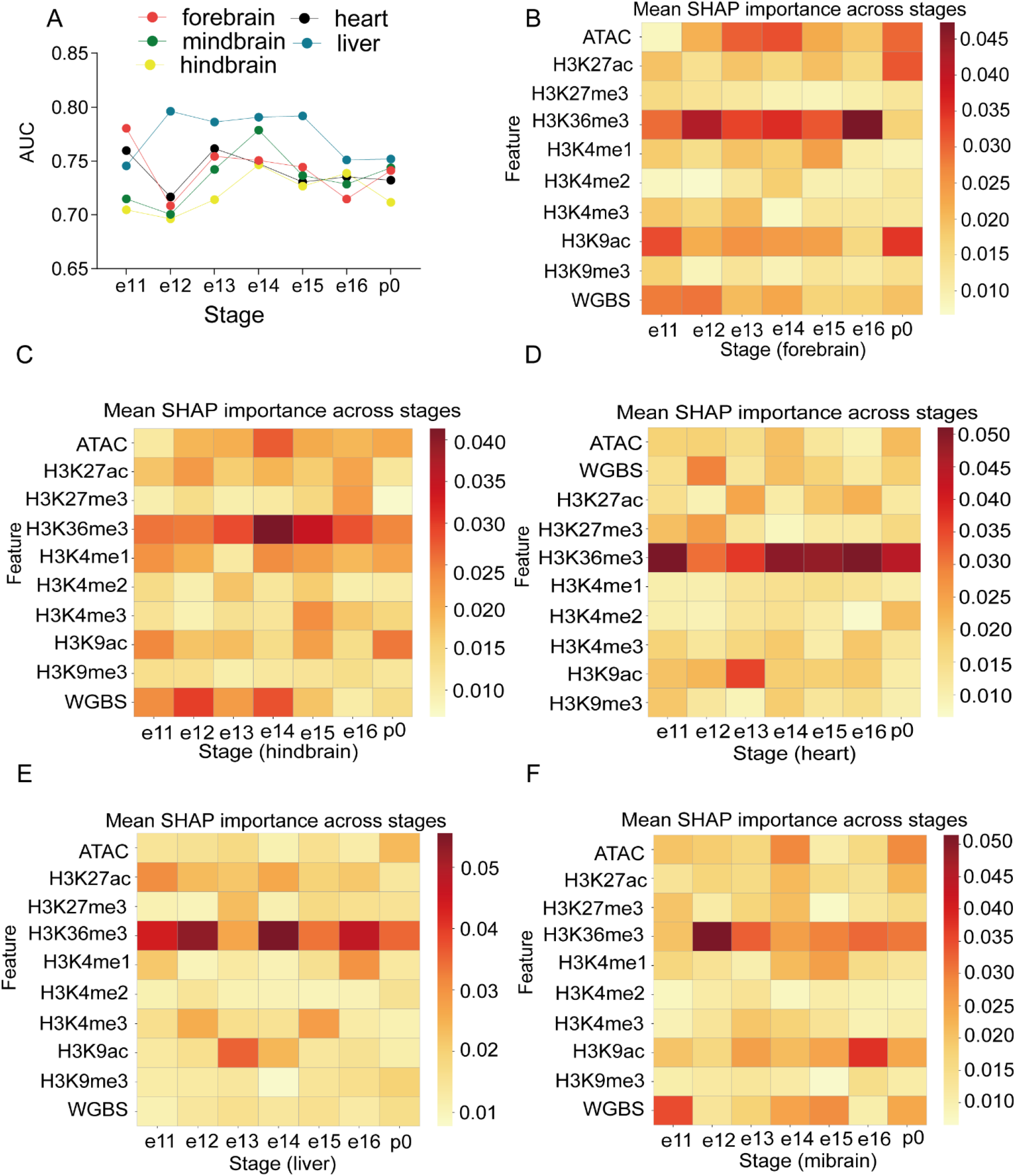
Epigenetic feature prediction of RNA editing frequency during mouse embryonic development. (A) AUC values of the Random Forest classifier for high-versus low-frequency editing sites across five tissues and seven developmental stages (e11–p0). (B-F) SHAP feature importance heatmaps for (B) forebrain, (C) hindbrain, (D) heart, (E) liver, and (F) midbrain. Color intensity represents the mean absolute SHAP value for each epigenetic feature (ATAC-seq, DNA methylation, H3K27ac, H3K27me3, H3K36me3, H3K4me1, H3K4me2, H3K4me3, H3K9ac, H3K9me3).

To identify the epigenetic determinants underlying model performance, we calculated the mean SHAP values of upstream and downstream signals for each editing site and compared feature importance across tissues and developmental stages. Among all examined features, H3K36me3 consistently ranked as the most influential predictor across virtually all datasets, whereas the contributions of other histone modifications, chromatin accessibility, and DNA methylation varied considerably among tissues and stages (Figure 7B–F). Notably, this dominant role of H3K36me3 was maintained despite marked species-specific differences in the genomic distribution of editing sites, with mouse editing events preferentially occurring in 3’ UTRs and SINE elements rather than the intronic and Alu repeats characteristic of the human genome.

The conserved predictive importance of H3K36me3 across diverse mouse tissues closely mirrors our findings in human cell lines, providing strong evidence that this histone modification represents a fundamental epigenetic correlate of A-to-I RNA editing frequency. The persistence of this association across multiple tissues, developmental stages, and two mammalian species suggests that H3K36me3-dependent chromatin organization constitutes a conserved regulatory framework for establishing RNA editing landscapes, whereas the contributions of other epigenetic features are likely to be more context dependent.

### Functional validation of H3K36me3 in the regulation of A-to-I RNA editing

To directly determine whether H3K36me3 regulates A-to-I RNA editing, we analyzed matched RNA-seq and ChIP-seq datasets generated from wild-type and SETD2-knockout renal epithelial cells, in which deposition of H3K36me3 is selectively abolished [55].

Because alterations in ADAR abundance could influence RNA editing, we first quantified the expression of the two major editing enzymes. Neither ADAR1 nor ADAR2 expression was significantly altered following SETD2 knockout (Figure 8A-B), indicating that subsequent changes in RNA editing are unlikely to result from altered ADAR expression and instead reflect chromatin-dependent regulation.

**Figure 8.**
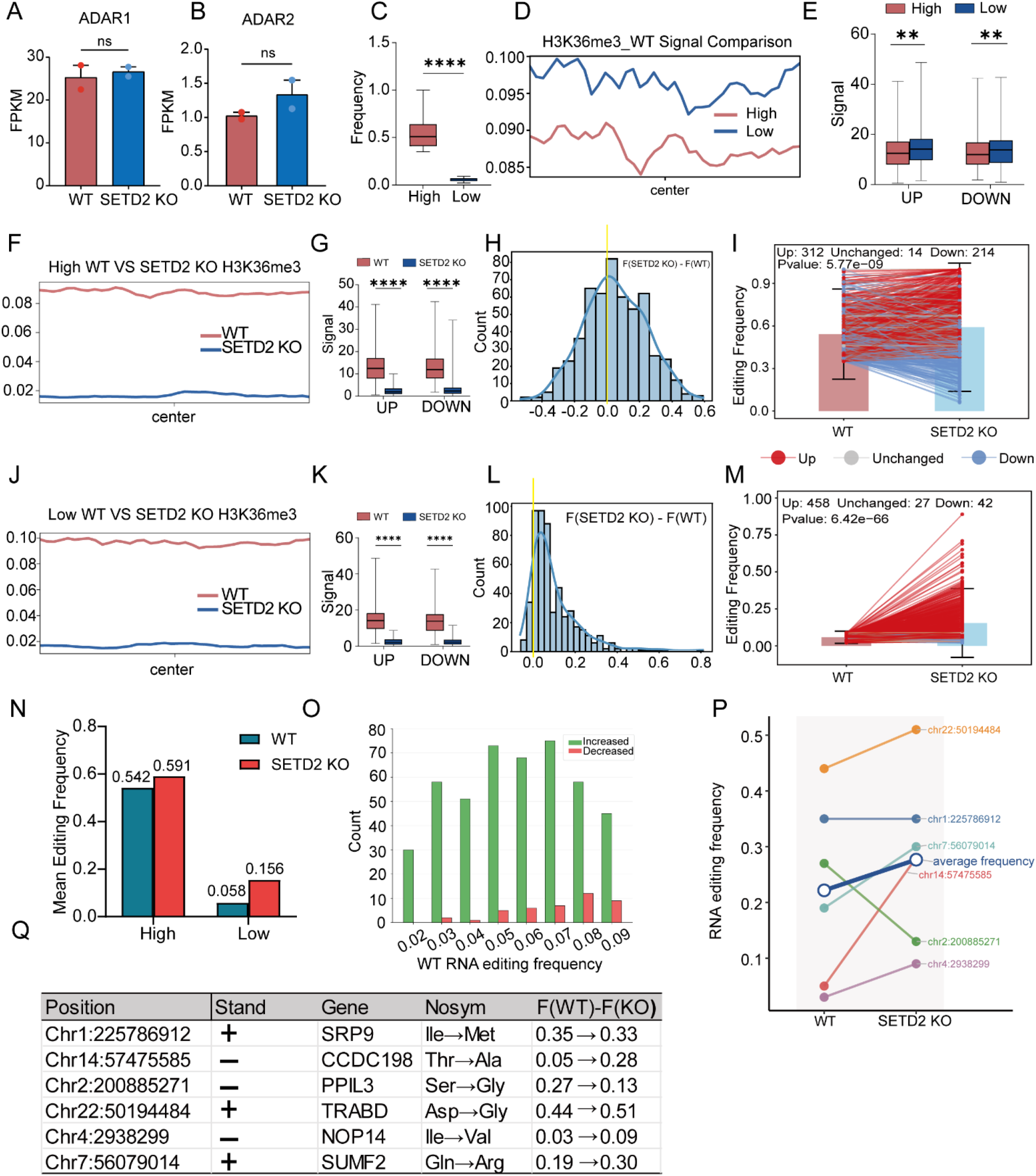
Loss of H3K36me3 upon SETD2 knockout derepresses A-to-I RNA editing. (A) ADAR1 transcript expression in wild-type (WT) and SETD2 knockout (KO) renal epithelial cells. (B) ADAR2 transcript expression in WT and SETD2 KO cells. (C) Editing frequency distribution in WT cells, stratified into high-frequency (≥75th percentile) and low-frequency (≤25th percentile) groups. (D) H3K36me3 ChIP-seq signal intensity within ±150 bp of high-frequency and low-frequency editing sites in WT cells. (E) Statistical comparison of H3K36me3 ChIP-seq signal intensity for upstream and downstream 150 bp of high-frequency versus low-frequency editing sites in WT cells. (F) H3K36me3 metagene profile at high-frequency editing sites in WT and SETD2 KO cells. (G) Statistical comparison of H3K36me3 ChIP-seq signal intensity for upstream and downstream 150 bp of high-frequency editing sites in WT cells versus in SETD2 KO cells. (H) Distribution of editing frequency changes at high-frequency editing sites following SETD2 KO. (I) Pairwise comparison of editing frequencies at high-frequency editing sites before and after SETD2 knockout. (J) H3K36me3 metagene profile at low-frequency editing sites in WT and SETD2 KO cells. (K) Statistical comparison of H3K36me3 ChIP-seq signal intensity for upstream and downstream 150 bp of low-frequency editing sites in WT cells versus in SETD2 KO cells. (L) Distribution of editing frequency changes at low-frequency editing sites following SETD2 KO. (M) Pairwise comparison of editing frequencies at low-frequency editing sites before and after SETD2 knockout. (N) Net change in mean editing frequency for high-and low-frequency groups upon SETD2 KO. (O) Relationship between baseline editing frequency in WT (x-axis) and the change in editing frequency following SETD2 knockout (KO − WT, y-axis) for low-frequency editing sites. (P) Editing frequency changes at individual nonsynonymous editing sites. (Q) Summary of nonsynonymous editing site responses upon SETD2 KO. **p < 0.01, ***p < 0.001, ****p < 0.0001; ns, not significant.

We next identified A-to-I RNA editing sites in WT and SETD2-knockout cells and compared editing frequencies between the two conditions. Editing sites were classified into high-and low-frequency groups based on the upper (≥75th percentile) and lower (≤25th percentile) quartiles of editing frequency in WT cells (Figure 8C). Consistent with our previous analyses, low-frequency editing sites exhibited significantly higher H3K36me3 enrichment within the ±150 bp regions flanking editing sites than high-frequency sites (Figure 8D-E), further supporting an inverse relationship between local H3K36me3 occupancy and RNA editing frequency.

As expected, deletion of SETD2 resulted in a marked global loss of H3K36me3 signal (Figure 8F and J). H3K36me3 levels surrounding editing sites were significantly reduced in both the high-and low-frequency groups relative to WT cells (Figure 8G and K), confirming effective disruption of local H3K36me3 deposition.

Loss of H3K36me3 was accompanied by a widespread increase in RNA editing frequency. High-frequency editing sites showed a modest overall increase, with a net gain of 0.049 following SETD2 knockout (Figure 8N). Although editing changes at these sites were bidirectional, 57.8% of sites exhibited increased editing compared with 39.6% showing decreased editing (Figure 8H and I). In contrast, low-frequency editing sites displayed a much stronger response to H3K36me3 depletion. Strikingly, low-frequency sites showed a ∼2.7-fold increase in editing frequency following SETD2 knockout (from 0.058 to 0.156), with 86.9% of sites upregulated (Figure 8L-N). This preferential derepression of low-frequency sites suggests that H3K36me3 normally acts as a ‘brake’ on suboptimal editing substrates. Notably, the magnitude of editing induction was inversely correlated with the baseline editing frequency in WT cells, with sites exhibiting the lowest initial editing levels showing the greatest increases. Indeed, virtually all editing sites with baseline frequencies ≤ 0.02 were upregulated following SETD2 depletion (Figure 8O). The inverse correlation between baseline editing frequency and the magnitude of induction following H3K36me3 loss suggests that sites with the strongest H3K36me3 occupancy are most repressed, consistent with a direct regulatory relationship.

To evaluate the functional consequences of these changes, we further examined nonsynonymous RNA editing events. Six nonsynonymous editing sites were identified across six genes. Four sites (TRABD, SUMF2, CCDC198, and NOP14) exhibited increased editing following SETD2 knockout, whereas two sites (SRP9 and PPIL3) showed modest decreases (Figure 8P and Q). Notably, consistent with the genome-wide analysis, nonsynonymous editing sites with low baseline editing frequencies displayed the largest increases following H3K36me3 depletion: from 0.05 to 0.28 for CCDC198 and from 0.03 to 0.09 for NOP14.

Collectively, these findings provide direct functional evidence that H3K36me3 restrains A-to-I RNA editing independently of changes in ADAR expression. Genetic ablation of SETD2, and the resulting loss of H3K36me3, preferentially enhanced editing at sites with low basal editing frequencies, establishing a causal link between local chromatin state and RNA editing efficiency. These results support a model in which H3K36me3 acts as an epigenetic barrier that limits RNA editing, thereby contributing to the establishment and maintenance of site-specific editing landscapes.

## DISCUSSION

A-to-I RNA editing is an essential post-transcriptional mechanism that expands transcriptomic diversity and regulates multiple aspects of RNA metabolism. Although ADAR1 and ADAR2 catalyze millions of editing events across the transcriptome, variation in ADAR expression explains only a fraction of the differences in editing levels observed across tissues, developmental stages, and disease states [21, 56], indicating the existence of additional regulatory mechanisms. While considerable progress has been made in understanding the catalytic functions of ADAR enzymes and the roles of RNA sequence, secondary structure, RNA-binding proteins, and RNA modifications in determining editing activity [22–31], the upstream mechanisms that determine why individual adenosines are edited at dramatically different efficiencies remain poorly understood. In this study, we identified local chromatin state as an important regulatory layer controlling RNA editing frequency. By integrating epigenomic profiling, machine learning, comparative analyses across human and mouse, and functional perturbation datasets, we identified H3K36me3 as a conserved negative regulator and H2A.Z.1 as a positive regulator of A-to-I RNA editing. These findings extend the functional repertoire of histone variants and modifications beyond transcriptional regulation, supporting a model in which chromatin architecture shapes the post-transcriptional RNA editing landscape.

A key contribution of this study is the demonstration that chromatin features serve as predictive determinants of RNA editing efficiency. Although previous studies have focused primarily on RNA-centered mechanisms, these factors alone do not fully explain the extensive variation in editing frequencies observed among genomic loci, tissues, developmental stages, and physiological conditions [21]. Our findings suggest that local chromatin organization establishes a permissive or restrictive environment that influences the probability of RNA editing before or during co-transcriptional RNA processing. This chromatin-centered perspective provides a broader framework for understanding how RNA editing is integrated into epigenetic regulation of gene expression.

Among all chromatin features examined, H3K36me3 emerged as the most robust and evolutionarily conserved predictor of RNA editing frequency. H3K36me3 is a well-established histone modification associated with transcription elongation and co-transcriptional RNA processing [57, 58]. Remarkably, this relationship was consistently observed across multiple human cell lines, five mouse tissues, and seven developmental stages despite substantial differences in genomic organization between species. Human RNA editing predominantly occurs within primate-specific Alu elements located in introns [43, 46], whereas mouse editing is largely associated with SINE elements and is enriched in 3′ UTRs. Nevertheless, H3K36me3 remained the dominant predictive feature in both species. This conservation suggests that chromatin-mediated regulation of RNA editing operates independently of the specific repetitive elements that generate editing substrates, and instead reflects a fundamental property of co-transcriptional RNA processing conserved throughout mammalian evolution.

Our functional analyses further demonstrated that the relationship between H3K36me3 and RNA editing is not merely correlative. Loss of H3K36me3 following SETD2 depletion resulted in widespread increases in RNA editing without detectable changes in ADAR1 or ADAR2 expression, indicating that chromatin can regulate editing independently of editing enzyme abundance. Upon SETD2 knockout, low-frequency editing sites exhibited a pronounced increase (86.9% of the sites upregulated with an approximately 2.7-fold increase in editing frequency), whereas highly edited sites displayed more heterogeneous responses (57.8% of the sites upregulated and 39.6% downregulated). The preferential increase in previously low-frequency editing sites further suggests that H3K36me3 normally suppresses editing at suboptimal substrates. These observations support a causal role for H3K36me3 in modulating editing efficiency and position chromatin state upstream of ADAR-mediated catalysis.

The molecular basis underlying this regulation remains to be elucidated, but several non-mutually exclusive models can be proposed. H3K36me3 is deposited during transcription elongation and is intimately associated with RNA polymerase II progression, nucleosome organization, and co-transcriptional RNA processing. Alterations in H3K36me3 may therefore influence transcriptional elongation kinetics, thereby modifying the temporal window during which nascent transcripts fold into double-stranded RNA structures accessible to ADAR enzymes. Alternatively, H3K36me3-dependent chromatin organization may regulate RNA polymerase II occupancy, nucleosome positioning, or higher-order chromatin architecture, indirectly affecting RNA folding and substrate accessibility. It is also conceivable that H3K36me3-associated chromatin complexes recruit as-yet-unidentified RNA-binding proteins or chromatin-associated factors that regulate ADAR localization or catalytic activity. Distinguishing among these possibilities will require future studies combining locus-specific chromatin editing, live-cell imaging of transcription, and analyses of ADAR-chromatin interactions.

In contrast to H3K36me3, the histone variant H2A.Z.1 emerged as a positive regulator of RNA editing. H2A.Z.1 has long been implicated in transcriptional activation, promoter accessibility, nucleosome dynamics, and transcriptional responsiveness [48, 51]. Consistent with these functions, depletion of H2A.Z.1 preferentially reduced editing at highly edited loci (76.3% of the sites decreased, mean Δ = −0.07), whereas low-frequency sites displayed heterogeneous responses with no overall directional change. These findings suggest that H2A.Z.1 is unlikely to function as a universal determinant of RNA editing but instead facilitates efficient editing within chromatin environments that are already permissive for ADAR activity. Together with the opposing effects of H3K36me3, these results demonstrate that distinct chromatin features can differentially modulate RNA editing efficiency, underscoring the complexity of chromatin-dependent regulation of post-transcriptional processes.

Increasing evidence suggests that RNA editing is tightly coupled to transcriptional and co-transcriptional RNA processing [39], both of which are strongly influenced by chromatin organization and epigenetic state. Our study identified H3K36me3 and H2A.Z.1 as prominent chromatin regulators with opposing effects on A-to-I RNA editing efficiency. By integrating computational prediction with functional perturbation analyses, we provided evidence that the epigenome predisposes certain genomic loci to differential RNA editing. These findings broaden our understanding of the epigenetic regulation of RNA editing and suggest that modulation of chromatin state may represent a potential strategy for correcting aberrant RNA editing in human disease. Future studies will be required to elucidate the molecular mechanisms linking chromatin architecture to ADAR recruitment and activity, as well as the cell-type-specific and disease-specific relevance of this regulatory pathway.

## MATERIALS and METHODS

### Data acquisition

To systematically investigate the landscape and epigenetic regulation of A-to-I RNA editing, we integrated matched transcriptomic and epigenomic datasets from both mouse and human sources [41, 42]. Mouse RNA-seq datasets from multiple tissues, including forebrain, midbrain, hindbrain, heart, and liver, were obtained from the ENCODE consortium and used for RNA editing identification and quantification. Corresponding WGBS, ATAC-seq, and ChIP-seq datasets profiling multiple histone modifications (H3K4me1, H3K4me2, H3K4me3, H3K9ac, H3K9me3, H3K27ac, H3K27me3, and H3K36me3) were collected to characterize chromatin accessibility and epigenetic features surrounding editing sites. For human analyses, matched RNA-seq and histone mark ChIP-seq datasets from three ENCODE cell lines (K562, GM12878, and HeLa-S3) were analyzed to evaluate the conservation of RNA editing landscapes and epigenetic regulation across species.

To functionally validate candidate epigenetic regulators, two independent GEO datasets were analyzed. The first (GSE134299) contains RNA-seq and H2A.Z.1 ChIP-seq data from wild-type and H2AFZ-deficient MCF-10A epithelial cells [54]. The second (GSE213260) includes RNA-seq and H3K36me3 ChIP-seq data from wild-type and SETD2-knockout renal epithelial cells, in which H3K36me3 deposition is specifically disrupted [55]. All datasets included at least two biological replicates. Together, these datasets provided a comprehensive multi-omics resource for investigating the epigenetic regulation of A-to-I RNA editing.

### RNA-seq processing and transcript quantification

RNA-seq datasets from mouse tissues and human cell lines were processed using a unified analysis pipeline. Raw sequencing reads were assessed using FastQC (v0.12.1), followed by adapter trimming and removal of low-quality bases (Phred score <20) using Trim Galore (v0.6.10). Reads shorter than 35 bp after trimming were discarded.

Filtered reads were aligned to the reference genome using STAR (v2.7.0f). Gene-level read counts were generated using HTSeq-count (v2.0.0), and expression levels were normalized as fragments per kilobase of transcript per million mapped reads (FPKM). Mouse reads were aligned to the GENCODE M25 (GRCm38) annotation, whereas human datasets were aligned to GENCODE Release 40 (GRCh38).

### Identification of A-to-I RNA editing sites

RNA editing sites were identified using REDItools (v1.3.1). Following quality control and STAR alignment, BAM files were sorted and indexed with SAMtools. Candidate editing sites were detected using REDItoolDnaRna.py based on RNA-reference mismatches and subsequently annotated using AnnotateTable.py with RepeatMasker and dbSNP annotations. Known genomic SNPs and mitochondrial editing sites were excluded to minimize false-positive calls. For mouse datasets, candidate sites were required to have a mapping quality ≥30, base quality ≥20, at least one edited read, and an editing frequency ≥2%. Only sites consistently detected across biological replicates were retained. For human datasets, region-specific filtering criteria were applied. Editing sites within Alu elements were required to have at least five supporting reads, one edited read, and an editing frequency ≥2%. Sites located in non-Alu repetitive or non-repetitive regions were filtered more stringently, requiring at least ten supporting reads, three edited reads, and an editing frequency ≥10%. To further eliminate mapping artifacts, candidate sites were subjected to pBLAT realignment. High-confidence editing sites were subsequently compared with REDIportal annotations and only reproducible sites identified across biological replicates were retained for downstream analyses.

### Motif analysis

To characterize sequence preferences surrounding A-to-I RNA editing sites, genomic sequences spanning two nucleotides upstream and downstream of each edited adenosine were extracted from the reference genome, generating 5-bp sequence windows centered on the editing site. For sites located on the negative strand, reverse-complement sequences were generated to maintain transcriptional orientation. Extracted sequences were exported in FASTA format and analyzed using MEME-ChIP to identify enriched sequence motifs and nucleotide composition surrounding editing sites.

### Functional annotation and enrichment analysis

RNA editing sites were functionally annotated using ANNOVAR to determine their genomic locations and associated genes. Gene Ontology (GO) enrichment analysis was performed using the R package clusterProfiler, evaluating enrichment in Biological Process, Molecular Function, and Cellular Component categories [59]. Significantly enriched GO terms were visualized using ggplot2.

### Epigenetic feature extraction

To characterize the chromatin environment surrounding RNA editing sites, ChIP-seq and ATAC-seq datasets were processed into normalized BigWig files using deepTools following BAM sorting and indexing with SAMtools. Average signal profiles surrounding editing sites were generated using deepTools, and quantitative signal intensities of histone marks and chromatin accessibility were extracted from BigWig files using pyBigWig. Unless otherwise specified, epigenetic signals were quantified within ±150 bp of each editing site.

### Machine learning analysis

RNA editing frequency was calculated as the proportion of edited reads among all reads covering each editing site. For datasets containing biological replicates, edited reads and total read counts were first summed across replicates before calculating editing frequencies, thereby minimizing variability arising from sequencing depth. Editing sites were ranked according to editing frequency and classified into high-frequency (≥75th percentile) and low-frequency (≤25th percentile) groups.

To investigate the contribution of epigenetic features to RNA editing frequency, machine learning and deep learning-based classifiers, including Logistic Regression (LR), Support Vector Machine (SVM), Random Forest (RF), eXtreme Gradient Boosting (XGBoost), Naïve Bayes, CatBoost, ResNet-style MLP, Extremely Randomized Trees (Extra Trees) classifier, and Extra Trees Full Ratio Regressor, were constructed using the extracted epigenetic feature matrix. Features were standardized prior to model training. Samples were randomly partitioned into five folds for cross-validation. During each iteration, four folds were used for training and the remaining fold for validation. Model performance was evaluated by the mean area under the receiver operating characteristic curve (AUROC) and precision-recall curve (AUPRC) across the five validation folds. Model interpretability was assessed using SHAP (SHapley Additive exPlanations), which quantifies the contribution of each feature to model predictions.

### Isotonic regression

The isotonic regression was implemented via the Pool Adjacent Violators Algorithm (PAVA). Briefly, observations were first sorted by the WT editing frequency. The algorithm initialized each observation as its own block and iteratively merged adjacent blocks that violated the monotonicity constraint. After each merge, the block mean was recomputed as a weighted average. This process continued until all block means were non-increasing, yielding a piecewise-constant isotonic fit.

### Statistical methods

Statistical analyses were performed using GraphPad Prism 10 and R. Two-way analysis of variance (ANOVA) was used for comparisons involving two independent variables, paired Student’s t-tests for comparisons between two paired groups, and Welch’s t-test for unpaired comparisons unless otherwise indicated. To control the false discovery rate (FDR), p-values were adjusted using the Benjamini–Hochberg procedure. Statistical significance was defined as an adjusted p-or q-value < 0.05, with significance levels indicated as *p < 0.05, **p < 0.01, ***p < 0.001, and ****p < 0.0001.

### Availability of software

The computational pipeline for A-to-I RNA editing analysis and accompanying online documentation are available at https://github.com/CAI9-728/A-to-I-RNA-Editing/.

## ACKNOWLEDGEMENTS

This work was supported by grants from the National Natural Science Foundation of China (No. 31771165, No.81971064 to Y. Z.), and funding from Wenzhou Medical University.

---

**Supplementary Figure 1.**
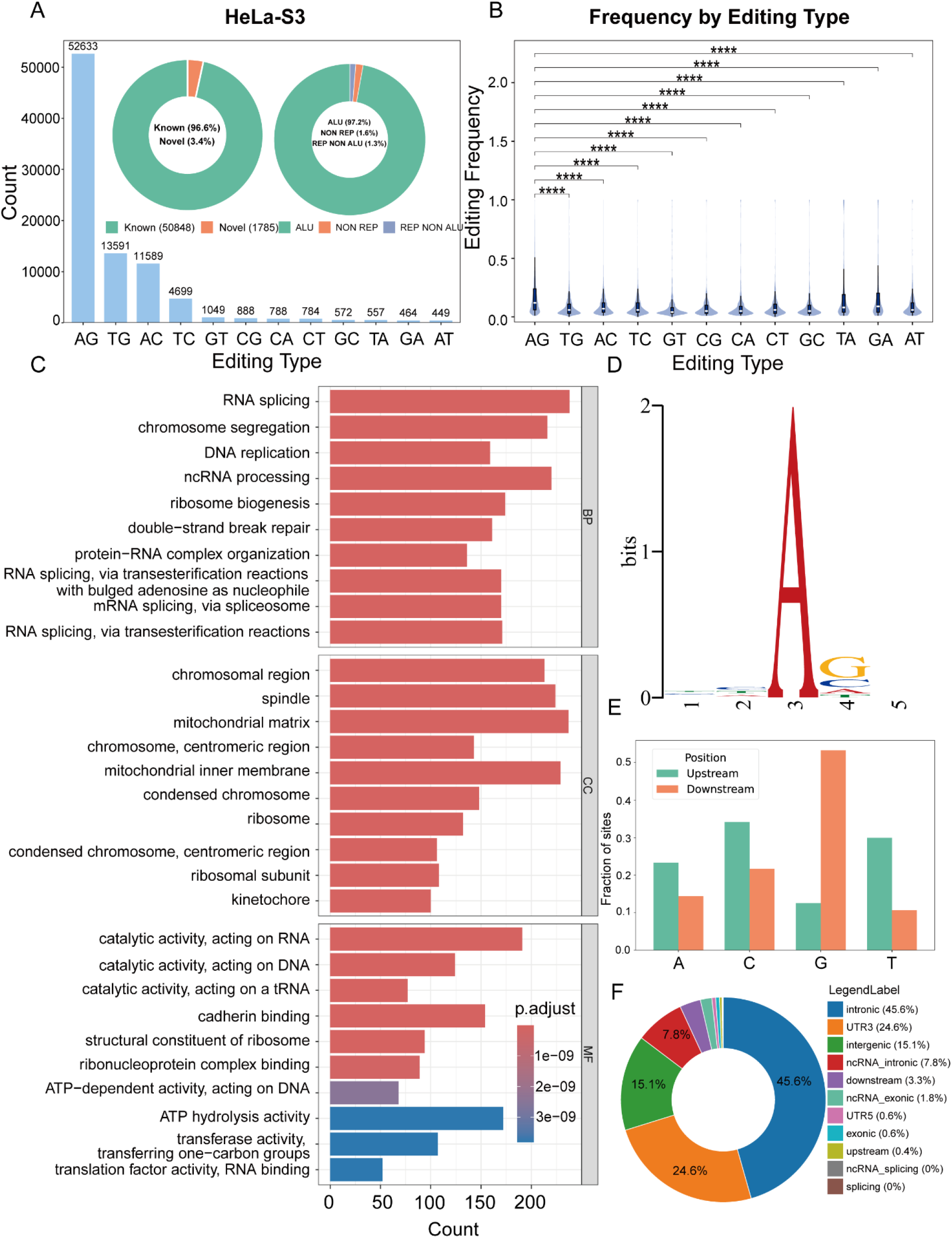
Identification and characterization of A-to-I RNA editing sites in HeLa-S3 cells. (A) Summary statistics of editing sites. (B) Relative frequencies of all 12 nucleotide substitution types, with A-to-G highlighted as the predominant editing signature. (C) Gene Ontology (GO) enrichment analysis of host genes harboring editing sites. BP, biological process; MF, molecular function; CC, cellular component. (D) Sequence logo showing the 2-bp nucleotide composition immediately upstream and downstream of editing sites. (E) Bar plot of 1-bp nucleotide frequencies at the −1 and +1 positions flanking editing sites. (F) Genomic distribution of editing sites. ****p < 0.0001.

**Supplementary Figure 2.**
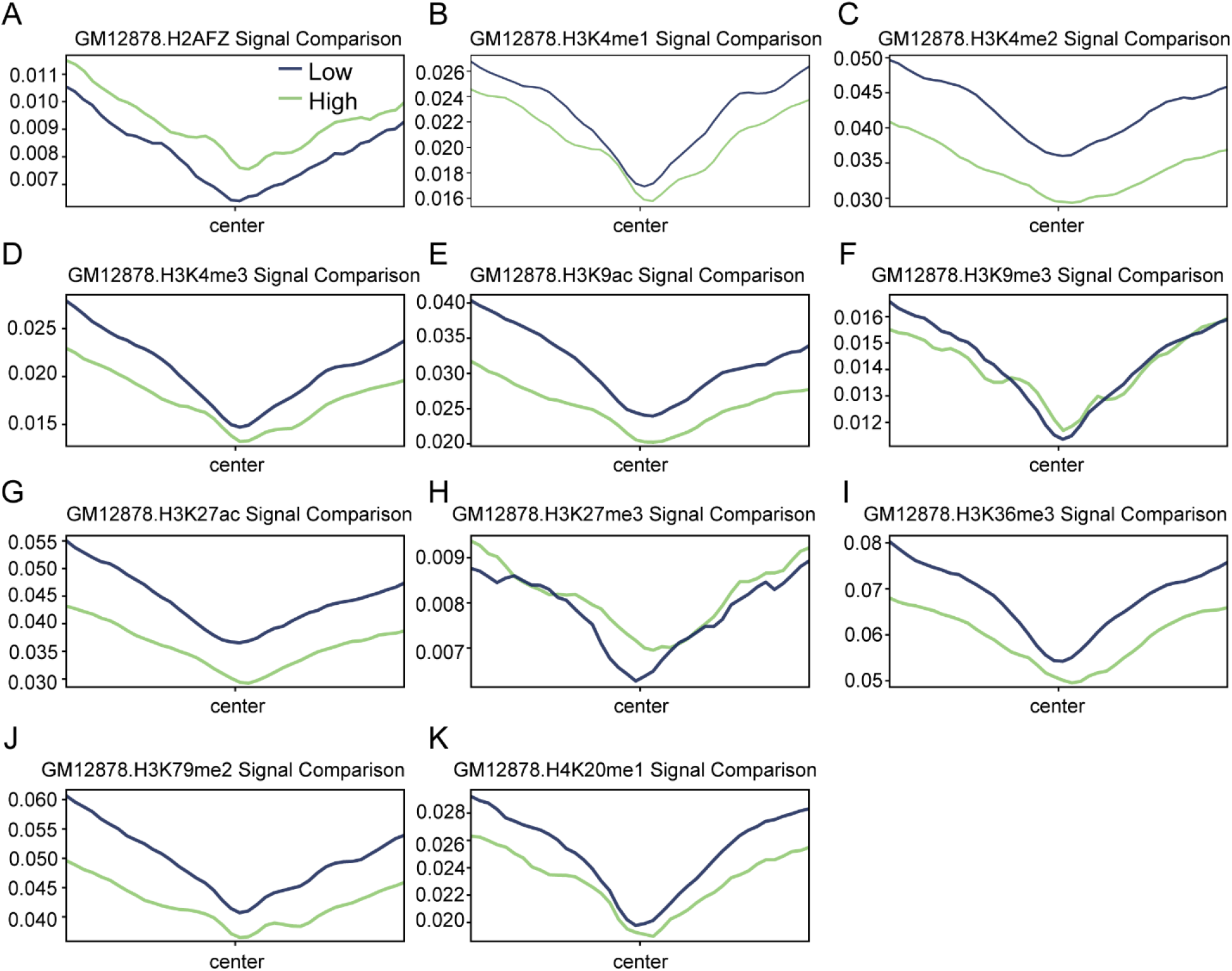
Histone feature profiles at high-and low-frequency A-to-I RNA editing sites in GM12878 cells. (A–K) Metagene plots showing average ChIP-seq signal intensity within ±150 bp of high-frequency (green; ≥75th percentile) and low-frequency (blue; ≤25th percentile) editing sites for (A) H2A.Z.1, (B) H3K4me1, (C) H3K4me2, (D) H3K4me3, (E) H3K9ac, (F) H3K9me3, (G) H3K27ac, (H) H3K27me3, (I) H3K36me3, (J) H3K79me2, and (K) H4K20me1.

**Supplementary Figure 3.**
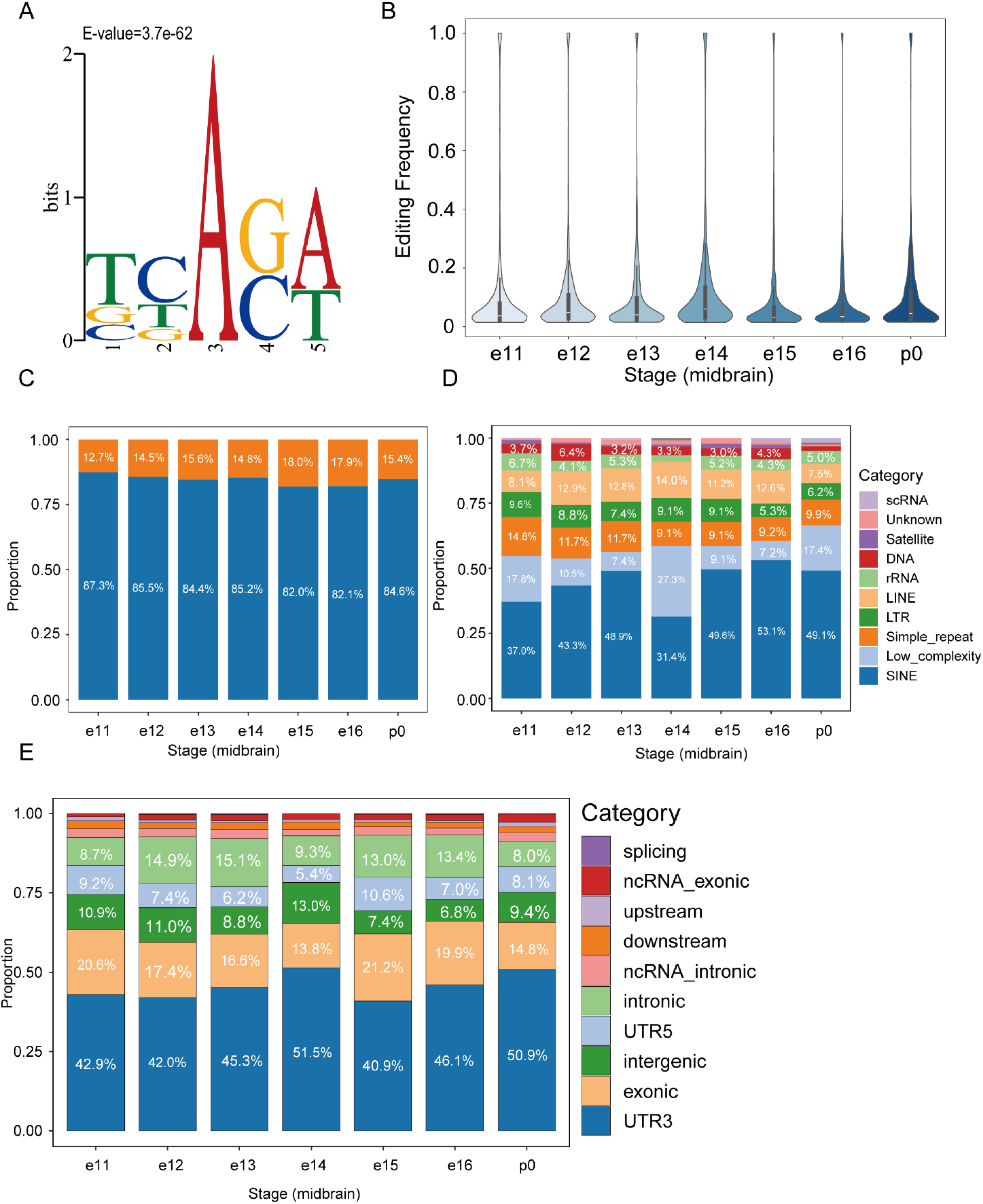
Distribution characteristics of A-to-I RNA editing sites in the mouse midbrain. (A) Sequence logo showing the 2-bp nucleotide composition upstream and downstream of editing sites. (B) Distribution of editing frequencies across embryonic development stages (e11–p0). (C) Proportion of editing sites in repetitive versus non-repetitive elements. (D) Genomic distribution of editing sites within repetitive elements. (E) Genomic distribution of editing sites.

**Supplementary Figure 4.**
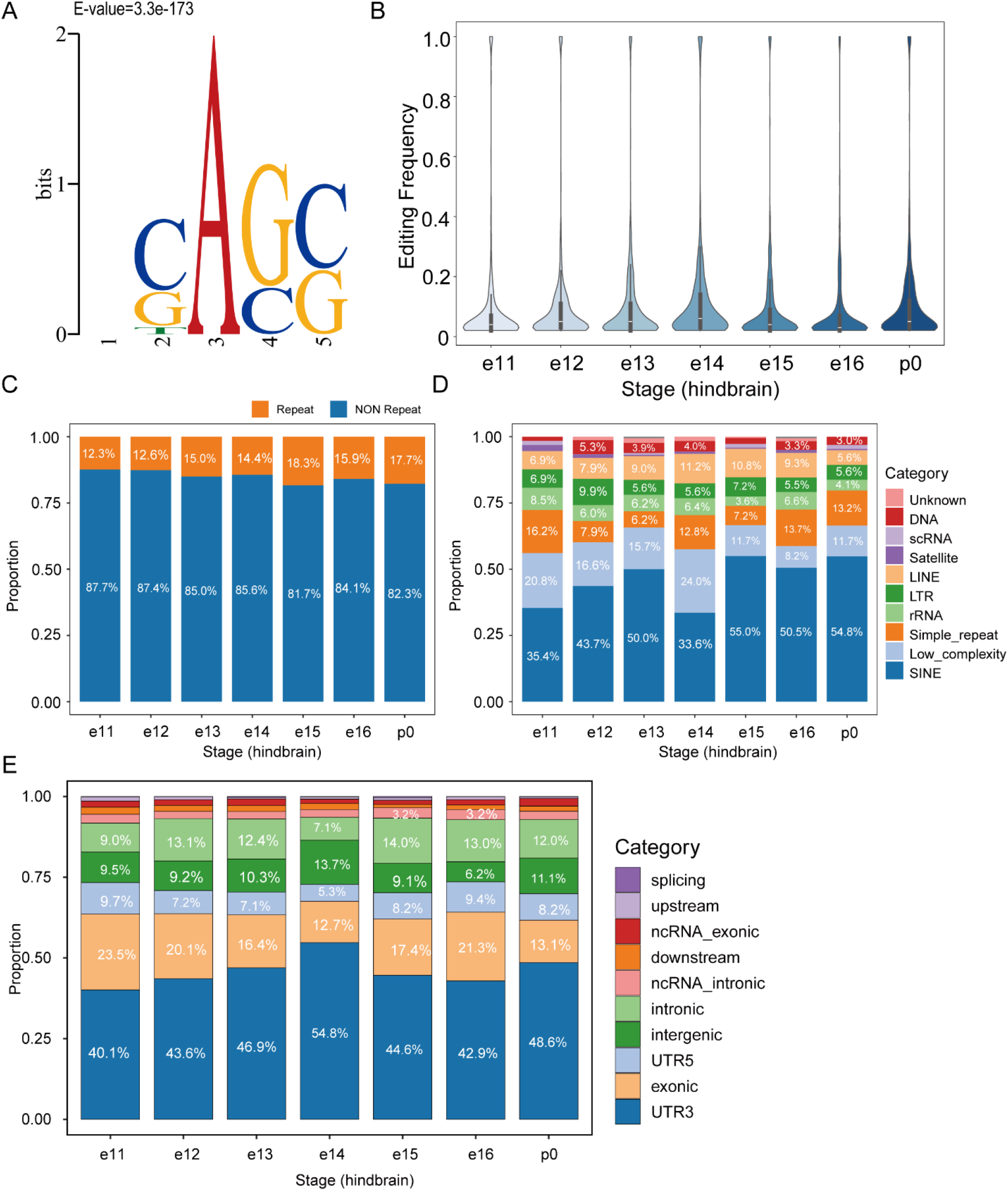
Distribution characteristics of A-to-I RNA editing sites in the mouse hindbrain. (A) Sequence logo showing the 2-bp nucleotide composition upstream and downstream of editing sites. (B) Distribution of editing frequencies across embryonic development stages (e11–p0). (C) Proportion of editing sites in repetitive versus non-repetitive elements. (D) Genomic distribution of editing sites within repetitive elements. (E) Genomic distribution of editing sites.

**Supplementary Figure 5.**
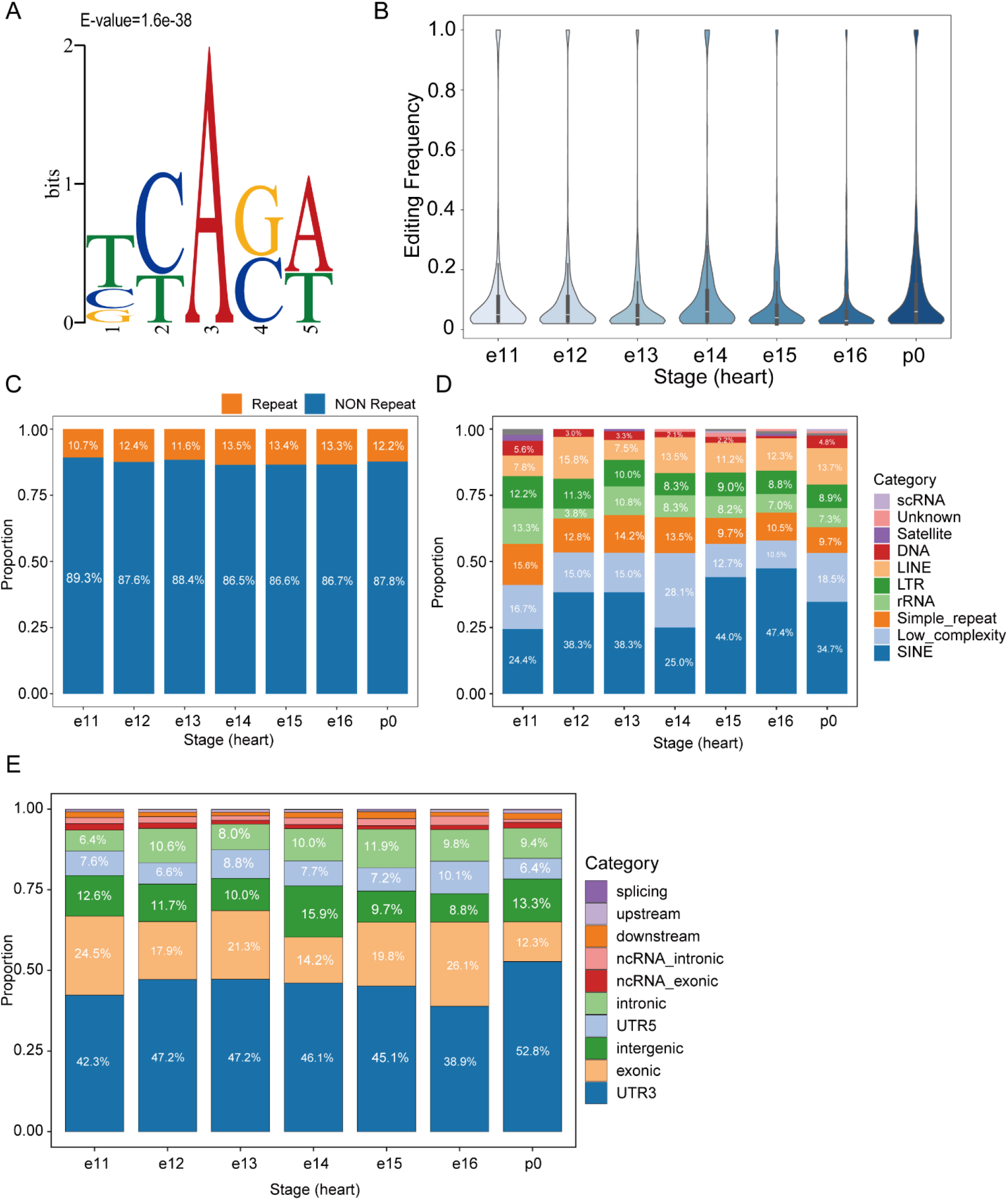
Distribution characteristics of A-to-I RNA editing sites in the mouse heart. (A) Sequence logo showing the 2-bp nucleotide composition upstream and downstream of editing sites. (B) Distribution of editing frequencies across embryonic development stages (e11–p0). (C) Proportion of editing sites in repetitive versus non-repetitive elements. (D) Genomic distribution of editing sites within repetitive elements. (E) Genomic distribution of editing sites.

**Supplementary Figure 6.**
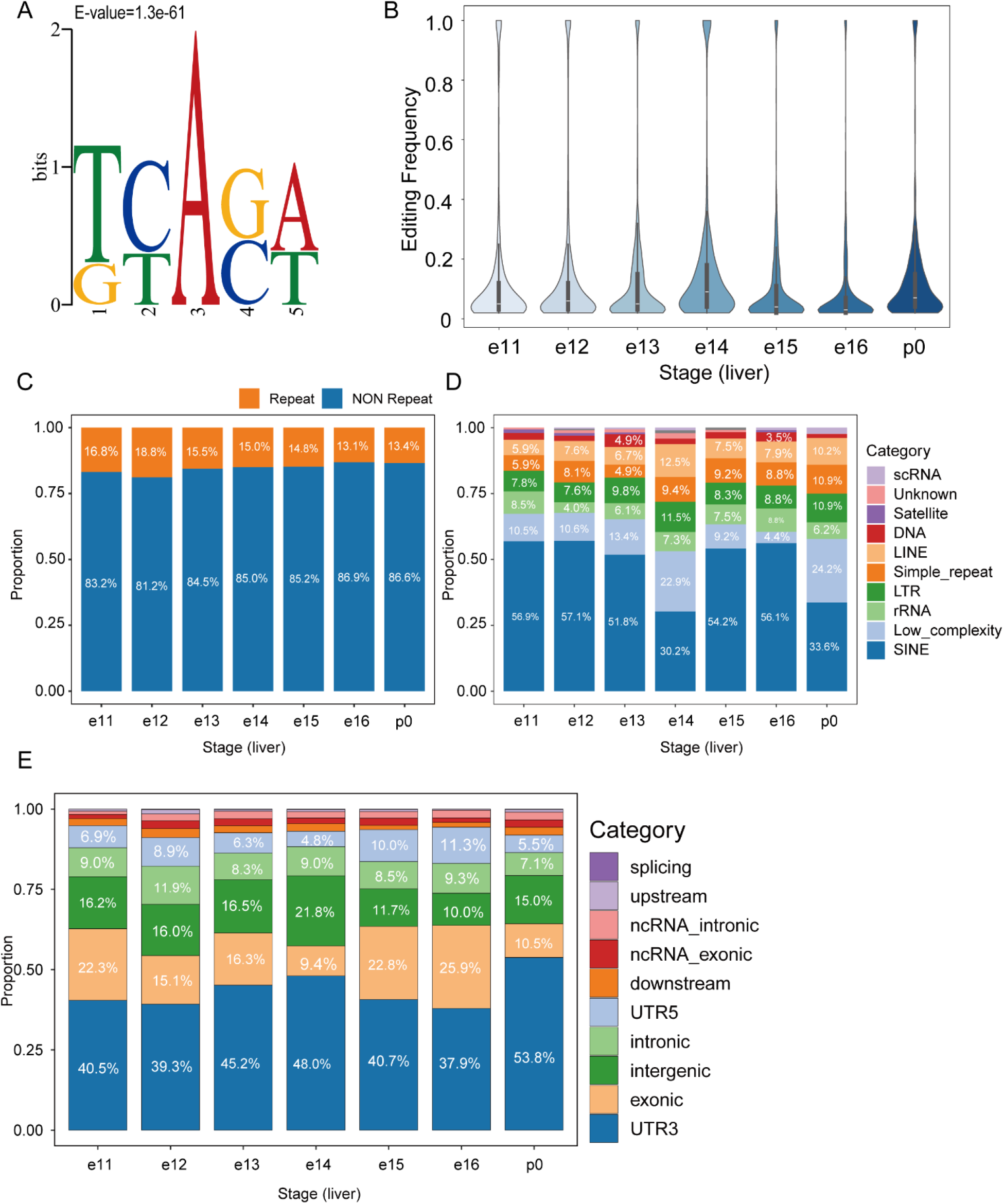
Distribution characteristics of A-to-I RNA editing sites in the mouse liver. (A) Sequence logo showing the 2-bp nucleotide composition upstream and downstream of editing sites. (B) Distribution of editing frequencies across embryonic development stages (e11–p0). (C) Proportion of editing sites in repetitive versus non-repetitive elements. (D) Genomic distribution of editing sites within repetitive elements. (E) Genomic distribution of editing sites.

**Supplementary Table 1.** Statistical comparison of ChIP-seq signals for each histone mark within ±150 bp of editing sites between high-and low-frequency editing groups in GM12878 cells, p-values were adjusted using the Benjamini–Hochberg (BH) procedure.

| Histone mark | High_frequency<br>mean | Low_frequency<br>mean | Fold change<br>(High/Low) | BH-FDR<br>q-value |
| --- | --- | --- | --- | --- |
| H3K36me3 | 17.07 | 19.50 | 0.88 | <1e-15 |
| H3K79me2 | 12.13 | 14.18 | 0.86 | 2.7e-14 |
| H3K4me3 | 4.94 | 5.80 | 0.85 | 4.3e-14 |
| H3K4me1 | 5.86 | 6.44 | 0.91 | 1.8e-10 |
| H3K4me2 | 9.76 | 12.24 | 0.80 | <1e-15 |
| H3K9ac | 7.20 | 8.86 | 0.81 | 1.2e-13 |
| H4K20me1 | 6.63 | 7.19 | 0.92 | 9.4e-15 |
| H3K27ac | 10.25 | 12.61 | 0.81 | 8.6e-11 |
| H2A.Z.1 | 2.63 | 2.30 | 1.15 | 7.4e-09 |
| H3K27me3 | 2.34 | 2.25 | 1.04 | 4.4e-03 |
| H3K9me3 | 4.05 | 4.09 | 0.99 | 0.567 |

